# Early megakaryocyte lineage-committed progenitors in adult mouse bone marrow

**DOI:** 10.1101/2021.08.06.455158

**Authors:** Zixian Liu, Jinhong Wang, Miner Xie, Peng Wu, Yao Ma, Sen Zhang, Xiaofang Wang, Fang Dong, Hui Cheng, Ping Zhu, Mingzhe Han, Hideo Ema

## Abstract

Hematopoietic stem cells (HSCs) have been considered to progressively lose their self-renewal and differentiation potentials prior to the commitment to each blood lineage. However, recent studies have suggested that megakaryocyte progenitors are generated at the level of HSCs. In this study, we newly identified early megakaryocyte lineage-committed progenitors (MgPs) in CD201^-^CD48^-^ cells and CD48^+^ cells separated from the CD150^+^CD34^-^Kit^+^Sca-1^+^Lin^-^ HSC population of the bone marrow in C57BL/6 mice. Single-cell transplantation and single-cell colony assay showed that MgPs, unlike platelet-biased HSCs, had little repopulating potential *in vivo*, but formed larger megakaryocyte colonies *in vitro* (on average eight megakaryocytes per colony) than did previously reported megakaryocyte progenitors (MkPs). Single-cell RNA-sequencing supported that these MgPs lie between HSCs and MkPs along the megakaryocyte differentiation pathway. Single-cell colony assay and single-cell RT-PCR analysis suggested the coexpression of *CD41* and *Pf4* is associated with megakaryocyte colony-forming activity. Single-cell colony assay of a small number of cells generated from single HSCs in culture suggested that MgPs are not direct progeny of HSCs. In this study, we propose a differentiation model in which HSCs give rise to MkPs through MgPs.

## Introduction

In the classical hierarchy model, hematopoietic stem cells (HSCs) progressively lose their self-renewal and differentiation potentials and then differentiate into unipotent progenitor cells ^1-4^. Recent studies using cell purification ^5-8^, cell tracking ^9^, and RNA sequencing technologies ^10,11^ have provided a different view of hematopoiesis. In particular, the megakaryocyte (Mk) lineage develops in close association with HSCs in the bone marrow of adult mice.

Mk differentiation potential has been detected in CD150^+^CD48^+^Flk2^-^c-Kit^+^Sca-1^+^Lin^-^ (KSL) cells (MPP2) ^12^, CD41^+^KSL cells ^13^, and CD150^+^CD41^+^c-Kit^+^Sca-1^-^Lin^-^ cells ^14^. Single-cell transplantation has revealed myeloid and Mk-lineage restricted repopulating progenitors in CD150^+^CD41^+^CD34^-^KSL cells ^5^. Platelet-biased (Plt-biased) HSCs, marked with vWF expression, have been reported as a subset of HSCs that give rise to only Mk lineage in primary transplantation but establish multilineage reconstitution in secondary transplantation ^8,9^. These HSCs seem to play a role under stressed conditions ^8,15^. It has been proposed that stem-like Mk committed progenitors (SL-MkPs) are primed to be of the Mk lineage by inflammation ^16^. Moreover, *in vivo* tracking studies have shown that Mks are an immediate progeny of HSCs in unperturbed hematopoiesis ^9^. Moreover, several studies have revealed similar gene expression profiles between HSCs and Mks, suggesting a developmentally close relationship ^7,9,12,17,18^. Some studies have suggested that megakaryocyte progenitors directly arise from HSCs without going through the stage of common myeloid progenitors (CMPs) or megakaryocyte/erythrocyte progenitors, both in mice and humans ^13,18,19^.

Despite all these studies, the relationship between HSCs and Mk-lineage committed progenitors, for instance, Mk differentiation pathways from HSCs, has not been thoroughly characterized to date. In this study, during a process to increase the purity of HSCs, we encountered new populations of early megakaryocyte progenitors (MgPs) in the mouse bone marrow. These MgPs appeared to be phenotypically similar to but functionally distinguishable from HSCs. Importantly, our identified MgPs showed little repopulating activity *in vivo*, suggesting that MgPs fundamentally differ from Plt-biased HSCs.

## Materials and methods

### Mice

C57BL/6 mice congenic for the Ly5 locus (B6-CD45.1) and C57BL/6 (B6-CD45.2) mice and β-actin-GFP transgenic B6 mice ^20^ were obtained from the animal facility of State Key Laboratory of Experimental Hematology. All the experimental protocols were approved by the Animal Care and Use Committee of the Institute of Hematology and Blood Disease Hospital.

### Flow-cytometric isolation

Bone marrow cells were obtained from 8- to 10-week-old B6 mice and stained with antibodies as described ^21^. Three lineage antibodies against Gr-1, B220, and Ter-119 (3Lin) were used ^22^. Cells were sorted into 96 well-plates using a BD FACSAria III. HSC1 was defined as the population of CD150^+^CD41^-^CD34^-^c-Kit^+^Sca-1^+^Lin^-^ cells. HSC2 was defined as the population of CD150^-^CD41^-^CD34^-^c-Kit^+^Sca-1^+^Lin^-^ cells. HPC1 was defined as the population of CD150^+^CD41^+^CD34^-^c-Kit^+^Sca-1^+^Lin^-^ cells (Figs. 1A-D). The HSC1, HSC2, and HPC1 (HSC1/HSC2/HPC1) populations were further divided into CD201^-^CD48^-^ cells (P1), CD201^+^CD48^-^ cells (P2); and CD48^+^ cells (P3) (Figs. 1E-G).

**Figure 1.**
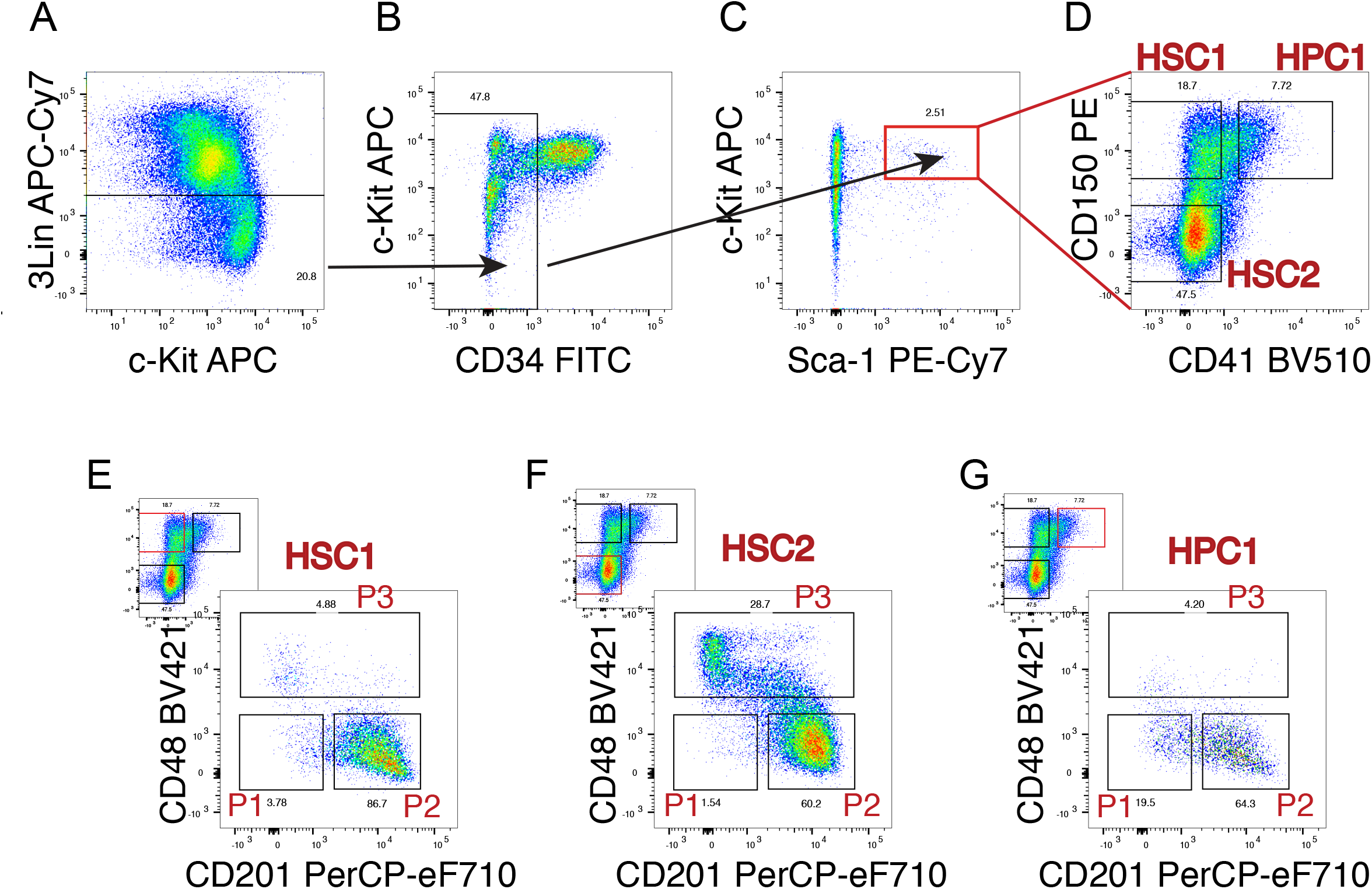
Gating strategy. Bone marrow cells were stained with antibodies against 3Lin, CD34, c-Kit, Sca-1, CD150, CD41, CD48, and CD201 antigens. (A) The gating of 3Lin^-^ cells. (B) The gating of CD34^-^ cells in 3Lin^-^ cells. (C) The gating of c-Kit and Sca-1 double-positive cells in CD34^-^Lin^-^ cells. (D) The gating of CD150^+^CD41^-^ cells (HSC1), CD150^+^CD41^-^ cells (HSC2), and CD150^+^CD41^+^ cells (HPC1) in CD34^-^KSL cells. (E) The HSC1 population was divided into 3 fractions (P1, 2, and 3) based on the expression of CD48 and CD201. (F) The HSC2 population was divided into 3 fractions (P1, 2, and 3) based on the expression of CD48 and CD201. (G) The HPC1 population was divided into 3 fractions (P1, 2, and 3) based on the expression of CD48 and CD201. P1, CD201^-^CD48^-^ cells; P2, CD201^+^ CD48^-^ cells; P3, CD48^+^ cells.

### Transplantation

Ten cells or single cells from bone marrow cells of B6-CD45.1 mice were sorted and transplanted into lethally irradiated B6-CD45.2 mice through the tail vein with 5×10^5^ bone marrow cells from B6-CD45.2 mice. In 10-cell transplantation, six months after the first transplantation, 2×10^7^ bone marrow cells were transplanted into 10 lethally irradiated B6-CD45.2 mice.

### Peripheral blood analysis

Peripheral blood of the recipient mice was analyzed by BD FACS Canto II for the myeloid, B cell, and T cell lineage contribution of donor-derived cells as described ^23^. To test the platelet (Plt) reconstitution in addition to three lineage reconstitution, GFP transgenic mice were used as donors. Successful reconstitution was defined as donor-derived cells in each lineage accounting for at least 0.05% at any time after transplantation. Long-term (LT)-HSCs were defined if the myeloid lineage reconstitution could be detected for at least 6 months after transplantation. Short-term (ST)-HSCs were defined if myeloid lineage reconstitution lasted less than 6 months after transplantation with B and T lymphoid reconstitution at any time after transplantation.

### Single-cell colony assay

Thirty single cells were sorted from P1, 2, and 3 fractions of HSC1, HSC2, and HPC1 populations (HSC1/HSC2/HPC1-P1/2/3), plated in an U-bottom 96-well plate by flow cytometry, and cultured in 200 µl α -MEM supplemented with 10% fetal bovine serum (FBS), 50 U of penicillin/streptomycin, 50 ng/ml murine stem cell factor (SCF), 50 ng/ml murine thrombopoietin (TPO), 10 ng/ml murine IL-3 (IL-3), and 1 IU/ml human erythropoietin (EPO). After 14 days of culture, differentiated cells were morphologically identified as neutrophils (n), macrophages (m), erythroblasts (E), and megakaryocytes (Mk) for each colony after Cytospin preparation. Mk colonies were identified by morphology under an inverted microscope, some of which were further stained with anti-CD41 antibody. On day 14, the diameters of nmEMk colonies were measured using ImageJ 1.52a software. Single cells were also isolated from CD150^+^CD48^+^Flk2^-^KSL cells (MPP2) and CD150^+^CD41^+^c-Kit^+^Sca-1^-^Lin^-^ cells (MkP population) and examined (Supplemental Fig. 1). Colonies are defined as those with ≥ 50 cells in one well. Besides, Mk colonies were defined as those with ≥ 3 cells in one well.

### Terminology

Early megakaryocyte progenitors MgPs were defined as megakaryocyte-lineage committed functional cells at the clonal level. Megakaryocyte progenitor MkP has been defined as the phenotypically defined population ^14^. Functional cells were designated as MkPs or MkP cells to distinguish them from the MkP population.

### Single-cell RNA sequencing (scRNA-seq)

HSC1/HSC2-P1/2/3 cells, as well as MPP2 cells were sorted into a 96-well plate with 10% FBS in PBS, and then for each population, 48 single cells were transferred by micromanipulator (Narishige, Japan) into 0.2-ml PCR tubes containing 2.55 μl lysis buffer with an 8-nt barcode. Single CD150^+^CD41^+^KSL cells (MkP) were directly sorted into 0.2 ml PCR tubes with lysis buffer. The scRNA-seq library was constructed with Smart-seq 2 (illumina) as previously reported ^24,25^. DNA sequencing was performed on an Illumina HiSeq 4000.

### Analysis of scRNA-seq data

After reads of adaptor contaminants and low-quality bases were removed, the reads were aligned to the mouse genome (GENCODE M16) using the STAR tool ^26^. The HTSeq package was used to count UMIs as the transcript copy number ^27^.

### UMAP plot and trajectory analysis

A total of 800 highly variable genes were used for the principal component analysis. Uniform manifold approximation and projection (UMAP) plot ^28^ was drawn using *RunUMAP* with dims set to 1:13. Monocle R package5 (v.2.18.0) was used to construct the trajectory tree of groups of cells, suggesting the differentiation orders ^29^.

### Identification of differentially expressed genes

We used the Seurat FindAllMarkers to identify unique cluster-specific marker genes ^30^. Gene ontology analysis was performed using DAVID (Bioinformatics Resources 6.8) ^31^.

### Single-cell reverse transcription polymerase chain reaction (single-cell RT-PCR)

Single cells from HSC1/HSC2/HPC1-P1/2/3 were sorted into PCR tubes containing 10 μL reverse-transcription and specific-target amplification mixture. RT-PCR was performed by Fluidigm Biomark as described ^23^. The sets of PCR primers, as listed in Supplemental Table 1, were purchased from Thermo Fisher Scientific.

### Statistical analyses

One-way ANOVA, Chi-square tests, Fisher’s exact test, and unpaired t test with Welch correction were performed with 23.0 (IBM). Wilcoxon test was performed with ggpubr R package (v.0.4.0). Significance was indicated as follows: *, p<0.05; **, p<0.01; ***, p<0.001; ****, p<0.0001.

## Results

### CD48 and CD201 divided HSC1, HSC2 and HPC1 into functionally distinct populations

The HSC1, HSC2, and HPC1 (HSC1/HSC2/HPC1) populations, respectively, were further divided into CD201^-^CD48^-^ (P1), CD201^+^CD48^-^ (P2) and CD48^+^ (P3) populations (Fig. 1). We performed transplantation with 10 P1, P2, and P3 cells each from the HSC1, HSC2, and HPC1 populations (HSC1/HSC2/HPC1-P1/2/3 cells). Fig. 2 shows the results of primary and secondary transplantation. Hematopoietic reconstitution was observed in 5 of 10 recipients of HSC1-P1 cells after primary transplantation but in none of the recipients after secondary transplantation (Fig. 2A). Reconstitution was observed in 8 of 10 recipients of HSC1-P2 cells after primary transplantation and in 8 of 10 recipients after secondary transplantation (Fig. 2A). Reconstitution was observed in only 1 of 10 recipients of HSC1-P3 cells after primary transplantation, the level of which decreased by 6 months and became undetectable after secondary transplantation (Fig. 2A).

**Figure 2.**
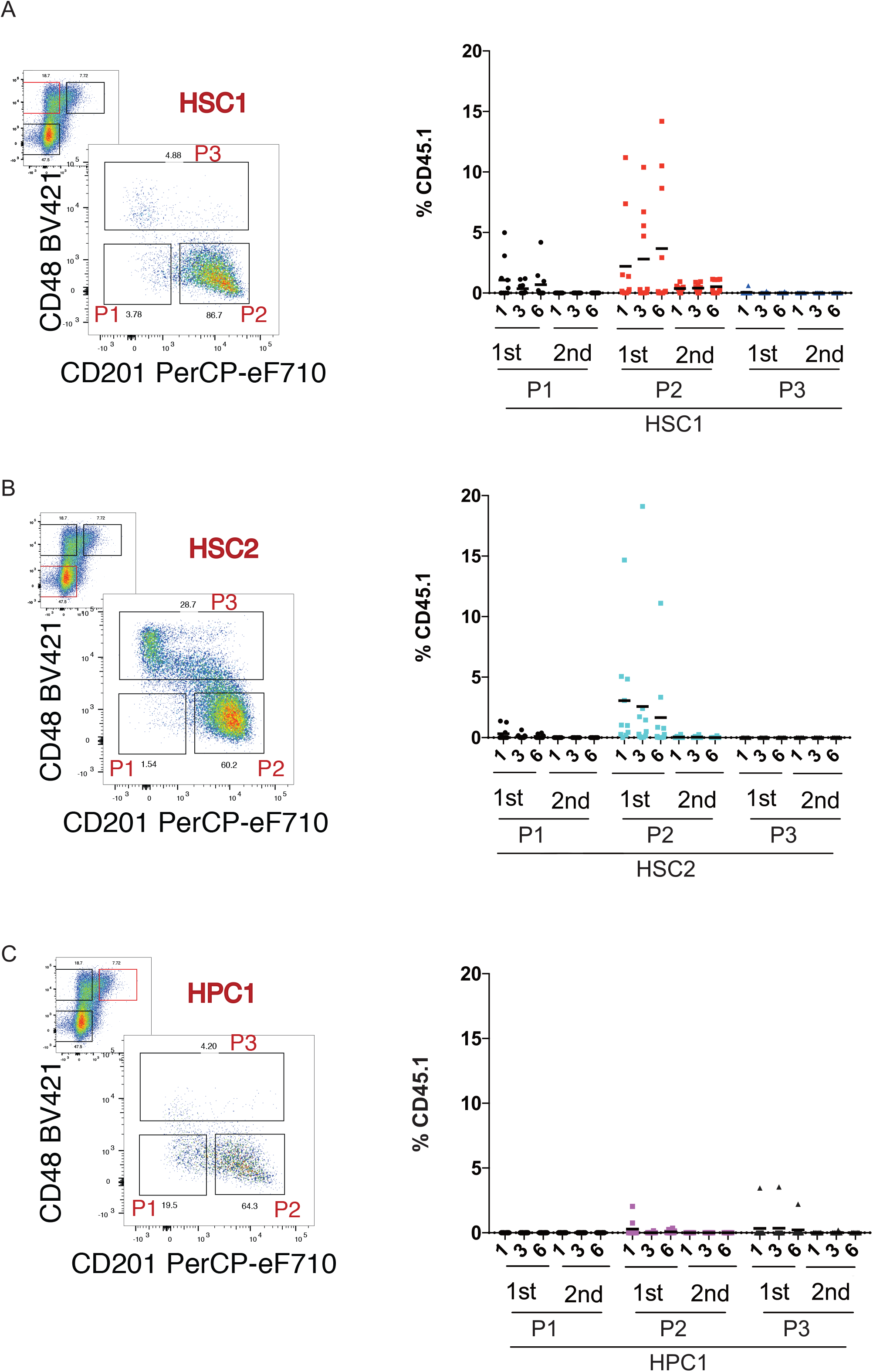
Long-term reconstitution with 10 cells. Ten cells each from the HSC1, HSC2 and HPC1 populations along with 5×10^5^ competitor cells were injected into 10 lethally irradiated mice. Peripheral blood was examined 1, 3, and 6 months after transplantation. Secondary transplantation was performed 6 months after transplantation. (A) Results of transplantation with 10 HSC1 cells. All mice survived. The reconstitution level 1 month after transplantation was similar between HSC1-P2 cells and HSC1-P1 cells (p=0.3718), but significantly greater in HSC1-P2 cells than in HSC1-P3 cells (p=0.0308). The reconstitution level 3 months after transplantation with HSC1-P2 cells was significantly greater than those with HSC1-P1 cells (p=0.0116) and HSC1-P3 cells (p=0.0032). The reconstitution level 6 months after transplantation with HSC1-P2 cells was significantly greater than those with HSC1-P1 cells (p=0.0015) and HSC1-P3 cells (p<0.0001). (B) Results of transplantation with 10 HSC2 cells. All mice survived. The reconstitution level 1 month after transplantation with HSC2-P2 cells was significantly greater than those with HSC2-P1 cells (p=0.0051) and HSC2-P3 cells (p=0.0015). The reconstitution level 3 months after transplantation with HSC2-P2 cells was significantly greater than those with HSC2-P1 cells (p=0.0131) and HSC2-P3 cells (p=0.0092). The reconstitution level 6 months after transplantation was similar between HSC2-P2 cells and HSC2-P1 cells or HSC2-P3 cells. (C) Results of transplantation with 10 HPC1 cells. Seven recipients of HPC1-P1 cells, 9 recipients of HPC1-P2 cells, and 7 recipients of HPC1-P3 cells survived. The reconstitution levels showed no significant difference at any time point after transplantation among HPC1-P1/2/3 cells.

Reconstitution was observed in 4 of 10 recipients of HSC2-P1 cells after primary transplantation but not after secondary transplantation. Reconstitution was observed in 9 of 10 recipients of HSC2-P2 cells after primary transplantation and in 2 of 10 recipients after secondary transplantation, but its level gradually decreased. No reconstitution was observed in recipients of HSC2-P3 cells after primary and secondary transplantation (Fig. 2B).

No reconstitution was observed in recipients of HPC1-P1 cells after primary and secondary transplantation. Reconstitution was observed in 5 of 10 recipients of HPC1-P2 cells after primary transplantation but not after secondary recipients. Reconstitution was observed in 1 of 10 recipients of HPC1-P3 cells after primary transplantation but not after secondary transplantation. The reconstitution level of HPC1-P2 cells was significantly lower and decreased faster than that of HSC1-P2 cells (Fig. 2C).

These results showed that the CD201^+^CD48^-^ populations (HSC1/HSC2/HPC1-P2) were enriched in reconstitution ability, while the CD201^-^CD48^-^ populations (HSC1/HSC2/HPC1-P1) showed lower and shorter reconstitution activity, and the CD48^+^ populations (HSC1/HSC2/HPC1-P3) showed little reconstitution activity.

### Detection of single repopulating cells

The frequencies of different types of repopulating cells were evaluated by single-cell transplantation. In this experiment, GFP transgenic mice were used as donors, and Plt reconstitution was also detected. The reconstitution rate of HSC1-P1 cells was 10.5% (2/19 mice) (Fig. 3A). All lineages, including myeloid, B cell, T cell and Plt lineages (My, B, T, P), were reconstituted in these two mice. However, the reconstitution disappeared by 6 months. By definition, these cells were short-term HSCs (ST-HSCs). The reconstitution rate of HSC1-P2 cells was 47.8% (11/23 mice). Six repopulating cells gave rise to all lineages (MyBTP), three of which were long-term (LT > 6 months)-HSCs, and the remaining three were ST-HSCs. Two repopulating cells gave rise to myeloid and lymphoid lineages (MyBT) and appeared to be ST-HSCs. One repopulating cell produced both B cells and Plts (BP). One cell was a B-lineage repopulating cell, and one was Plt-restricted repopulating cell (Plt-RC). No reconstitution was found in mice that received HSC1-P3 cells (Fig. 3A).

**Figure 3.**
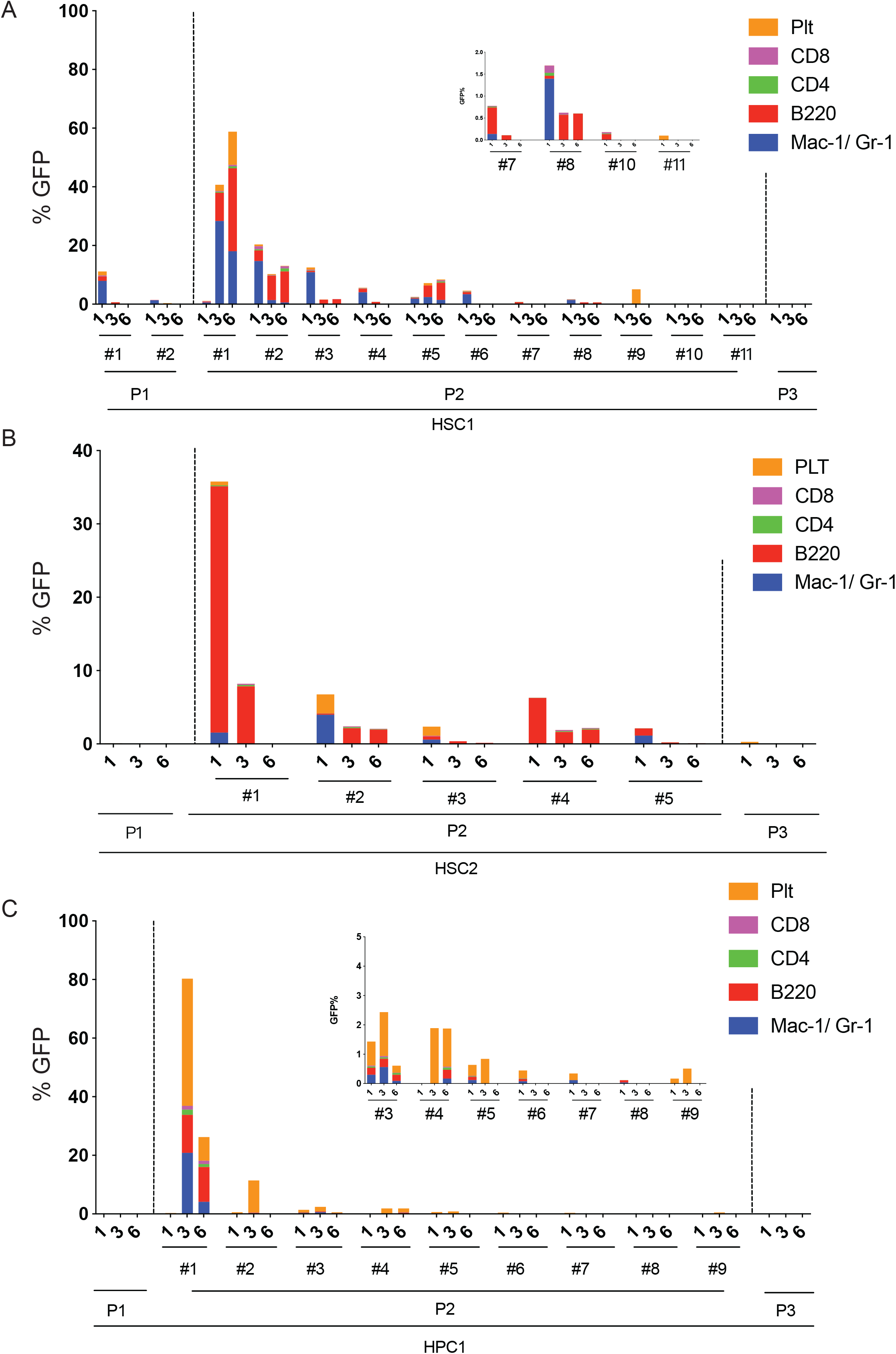
Single-cell transplantation. Single cells were isolated from (A) HSC1-P1/2/3, (B) HSC2-P1/2/3, and (C) HPC1-P1/2/3 populations, mixed with 5×10^5^ competitor cells, and injected into 30 lethally irradiated mice. Recipients were analyzed for 6 months after transplantation. Data of reconstituted mice are selectively shown (mouse ID is given by # number). Each column represents the percentage of total chimerism composed of myeloid, B cell, CD4 T cell, CD8 T cell, and Plt lineages. Data of mice with low chimerism are shown in the windows with enlargement. (A) Nineteen recipients of HSC1-P1 cells, 23 recipients of HSC1-P2 cells, and 20 recipients of HSC1-P3 cells survived. (B) Nineteen recipients of HSC2-P1 cells, 16 recipients of HSC2-P2 cells, and 12 recipients of HSC2-P3 cells survived. (C) Fourteen recipients of HPC1-P1 cells, 18 recipients of HPC1-P2 cells, and 14 recipients of HPC1-P3 cells survived. % GFP^+^ leukocytes and % GFP^+^ Plts are together shown in the total of 200 % (100% leukocytes and 100% Plts).

No reconstitution was found in recipients receiving HSC2-P1 cells (Fig. 3B). The reconstitution rate of HSC2-P2 cells was 31.25% (5/16). Two repopulating cells gave rise to all lineages (MyBTP) and appeared to be ST-HSCs. Two repopulating cells gave rise to myeloid and lymphoid lineages (MyBT), and one repopulating cell gave rise to myeloid and B lymphoid lineages (MyBT), all of which appeared to be ST-HSCs. One was a myeloid progenitor. Only one of the recipients of HSC2-P3 cells showed Plt reconstitution (Fig. 3B).

No reconstitution was found in recipients of HPC1-P1 cells (Fig. 3C). The reconstitution rate of HPC1-P2 cells was 50% (9/18). Three repopulating cells gave rise to all lineages (MyBTP) and turned out to be LT-HSCs. Three repopulating cells gave rise to all lineages (MyBTP) and turned out to be ST-HSCs. One repopulating cell gave rise to myeloid and lymphoid lineages and turned out to be ST-HSCs. One repopulating cell gave rise to transient reconstitution of myeloid and Plt (MyP) and turned out to be a myeloid progenitor. One repopulating cell was Plt-RC. No reconstitution was found in recipients of HPC1-P3 cells (Fig. 3C).

These single-cell transplantation data showed that HSC1/HSC2/HPC1-P2 cells were able to reconstitute multilineages. LT-HSCs were found in HSC1/HPC1-P2 cells. The reconstitution of these cells was characterized by myeloid-biased reconstitution at the beginning and gradually changed to balanced reconstitution. Most repopulating cells in HSC2-P2 cells appeared to be lymphoid-biased ST-HSCs. Plt-RCs were detected in HSC1-P2, HSC2-P3, and HPC1-P2 cells at a low frequency.

### nmEMk and Mk colony-formation

To further characterize the HSC1/HSC2/HPC1-P1/2/3 populations, we performed single-cell colony assay. Two kinds of colonies were observed. One was regular colonies (Fig. 4A, left panel). The other was Mk colonies comprising a small number of large cells expressing CD41, as detected by immunofluorescent staining (Fig. 4A middle and right panel). Colony cells were morphologically classified into neutrophils (n), macrophages (m), erythroblasts (E), and megakaryocytes (Mk) after Cytospin preparation of individual colonies (Supplemental Fig. 2A). Multilineage colonies with nmEMk, nmMk, or nmE lineages, bipotent colonies with nm lineages, and unipotent colonies with n, m, or Mk lineage were identified (Supplemental Fig. 2B). The colony-forming efficiencies were similar among the HSC1/HSC2/HPC1-P1/2/3 populations (Fig. 4B).

**Figure 4.**
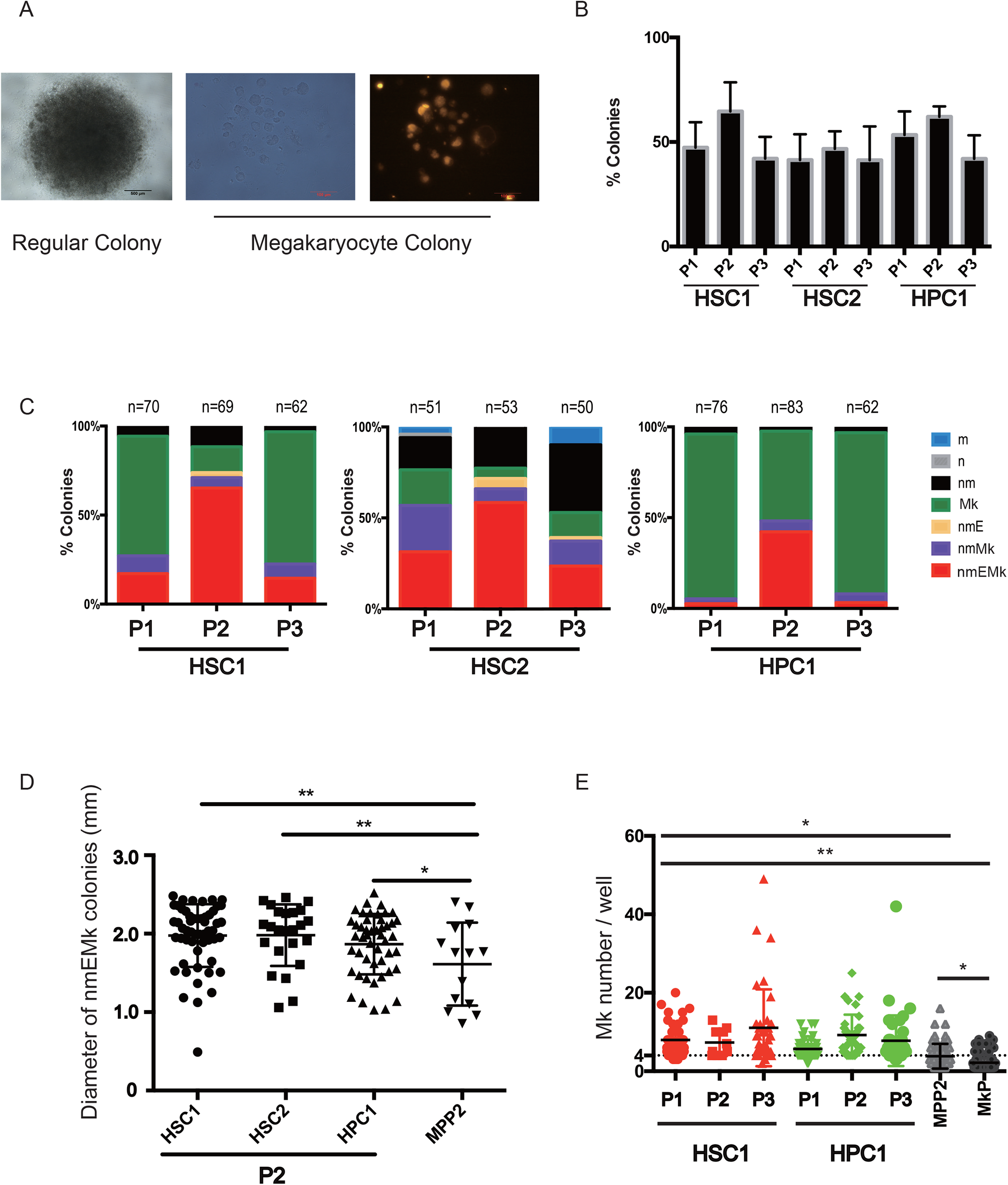
Single-cell colony assays. (A) Two types of colonies on day 14 of culture. Single cells were cultured in liquid medium with an U-bottom 96-well plate. The left panel shows a regular colony. The middle panel shows an Mk colony. The right panel shows the expression of CD41 (the same colony in the middle and right panels). (B) The frequencies of colony-forming cells. (C) Colony types identified. The frequency of nmEMk colony-forming cells in HSC1-P2 cells was significantly greater than those in HSC1-P1 cells (65.2% versus 17.1%, p < 0.001) and HSC1-P3 cells (65.2% versus 14.5%, p < 0.001); that in HSC2-P2 cells than those in HSC2-P1 cells (58.5% versus 31.4%, p = 0.001) and HSC2-P3 cells (58.5% versus 24%, p < 0.001); that in HPC1-P2 cells than those in HPC1-P1 cells (42.2% versus 2.6%, p < 0.001) and HPC1-P3 cells (42.2% versus 3.2%, P < 0.001) by chi-square test. The frequency of Mk colony-forming cells in HSC1-P2 cells was significantly smaller than those in HSC1-P1 cells (14.5% versus 67.1%, p < 0.001) and HSC1-P3 cells (14.5% versus 74.2%, p < 0.001); that in HPC1-P2 cells than those in HPC1-P1 cells (49.4% versus 90.8%, p < 0.001) and HPC1-P3 cells (49.4% versus 88.7%, p < 0.001) by chi-square test. Taken into account the colony-forming efficiency, Mk colonies were formed by 31.3 ± 10.4 % of HSC1-P1 cells, 6.7 ± 7.8 % of HSC1-P1 cells, 30.7 ± 13.8 % of HSC1-P3 cells, 6.7 ± 4.1 % of HSC2-P1 cells, 2.0 ± 3.0 % of HSC1-P2 cells, 4.0 ± 4.3 % of HSC2-P3 cells, 46.0 ± 13.0 % of HPC1-P1 cells, 27.3 ± 11.9 % of HPC1-P2 cells, and 36.7 ± 8.5 % of HPC1-P3 cells (mean ± S.D., n=5). n, neutrophil; m, macrophage; E, erythroblast; and Mk, megakaryocyte. (D) Day 14 nmEMk colony size. The diameters of nmEMk colonies were measured using ImageJ 1.52a software. nmEMk colonies formed by HSC1/HSC2/HPC1-P2 cells were significantly larger than those formed by MPP2 cells (p=0.003, p=0.008, and p=0.042, respectively, one-way ANOVA test). (E) Day 14 Mk colony size. A significant proportion of MPP2 and MkP cells did not divide, but matured into Mks. These cells were included in the size comparison. The numbers of Mks per colony were 8.0 ± 6.5 for HSC1/HPC1-P1/3 cells (mean ± S.D., n=167), 3.8 ± 3.2 for MPP2 cells (n=58), and 2.2 ± 2.0 for MkP cells (n=87). The number of Mks per colony of MPP2 cells was significantly smaller than those of HSC1-P1/2/3 cells (p<0.001, p=0.038, and p<0.001) and those of HPC1-P1/2/3 cells (p=0.047, p < 0.001, and p < 0.001; one-way ANOVA test). The number of Mks per colony of MkP cells was significantly smaller than those of HSC1-P1/2/3 cells (P<0.001, p=0.002, and p < 0.001) and those of HPC1-P1/2/3 cells (p<0.001 each; one-way ANOVA test). The number of Mks per colony of MkP cells was significantly smaller than that of MPP2 cells (p = 0.044; one-way ANOVA test).

The distribution of nmEMk colonies among the P1/2/3 populations was similar among the HSC1/HSC2/HPC1 populations: the highest frequency of nmEMk colonies was found in P2, and the proportion of nmEMk colonies was similar between P1 and P3. This finding suggested the enrichment of multipotent cells in P2 in common with the HSC1/HSC2/HPC1 populations (Fig. 4C).

There was no significant difference in the diameters among nmEMk colonies derived from the HSC1/HSC2/HPC1-P2 populations. MPP2 cells also formed nmEMk colonies (Supplemental Fig. 3A). The diameters of nmEMk colonies derived from the HSC1/HSC2/HPC1-P2 populations were significantly greater than those from the MPP2 population (Fig. 4D).

Mk colonies were formed by all HSC1/HSC2/HSC3-P1/2/3 populations. The majority (92.5 ± 26.8 %, n=5) of these Mk colonies were formed by the HSC1 and HPC1 populations (Fig. 4C). A large proportion (73.3 ± 31.3 %, n=5) of these Mk colonies was formed by the HSC1/HPC1-P1/3 populations. No difference was seen in the numbers of Mks per colony among the HSC1/HPC1-P1/3 populations. We also detected Mk colony-formation by single MPP2 cells (CD150^+^CD48^+^Flk2^-^KSL cells) and MkP cells (CD150^+^CD41^+^c-Kit^+^Sca-1^-^Lin^-^ cells) (Supplemental Fig. 3A). The numbers of Mks per colony (8.0 ± 6.5, n=167) in the HSC1/HPC1-P1/3 populations were significantly greater than those in the MPP2 and MkP populations (Fig. 4E). A large proportion of MkP cells generated only 1-3 Mks (Supplemental Fig. 3A), consistent with previous studies ^9,14^. The number of Mks per colony in the MPP2 population was significantly greater than that in the MkP population (Fig. 4E).

On day 14, some colonies were too small for Cytospin preparation for morphological analysis. To address this problem, we extended the culture period to 3 weeks. Some small colonies became larger by day 21 (Supplemental Fig. 3B). The lineage output of those slowly growing colonies appeared to be consistent with that of day 14 colonies. The percentages of nmEMk colonies in the HSC1/HSC2/HPC1-P2 populations were significantly greater than those in the HSC1/HSC2/HPC1-P1/3 populations. The percentages of Mk colonies in the HSC1/HPC1-P2 populations were significantly lower than those in the HSC1/HPC1-P1/3 populations.

In order to examine the relationship between nmEMk and Mk colony-forming cells, we performed single-cell colony assay on paired-daughter cells derived from single HSC1-P2 and HPC1-P2 cells. As shown in Supplemental Table 2, only one of 22 pairs derived from single HSC1/P2 formed the combination of nmEMk and Mk colonies, and none of 28 pairs derived from single HPC1/P2 cells formed such a combination, consistent with our previous study ^32^. A substantial number of nm/m colony-forming cells, but few Mk colony-forming cells appeared after the first division.

To further investigate myeloid lineage commitment in early divisions of HSCs, when single HSC1-P2 gave rise to 3-8 cells in culture, individual cells were isolated by micromanipulation and their colony forming potential was examined. As shown in Supplemental Table 3, a total of 224 cells derived from 43 single cells were analyzed. >50% of the cells formed nmEMk colonies, showing the amplification of nmEMk colony-forming cells in early divisions. >20% of the cells formed nm or m (nm/m) colonies, suggesting EMk or nEMk differentiation potential was lost after 2 or 3 division. Among them, however, Mk colonies were extremely rare. These data suggested that cells with nmEMk differentiation potential were committed to nm/m lineage from the first division, but the commitment to Mk lineage required 4 or more divisions.

### Relationship among HSCs and HPCs predicted by single-cell RNA-seq data

We performed single-cell RNA-sequencing (scRNA-seq) analysis on the HSC1/HPC1-P1/2/3, MPP2, and MkP populations to deduce the relationship among these populations. scRNA-Seq was successfully performed on 48 HSC1-P1 cells, 48 HSC1-P2 cells, 44 HSC1-P3 cells, 44 HPC1-P1 cells, 48 HPC1-P2 cells, 44 HPC1-P3 cells, 48 MPP2 cells, and 85 MkP cells. Transplantation and colony assay data suggested that the HSC1-P2 population was closely related with the HPC1-P2 population so that these populations were analyzed together, and that the HSC1-P1/3 populations were closely related with the HPC1-P1/3 populations so that these populations were analyzed together, as shown in Fig. 5.

**Figure 5.**
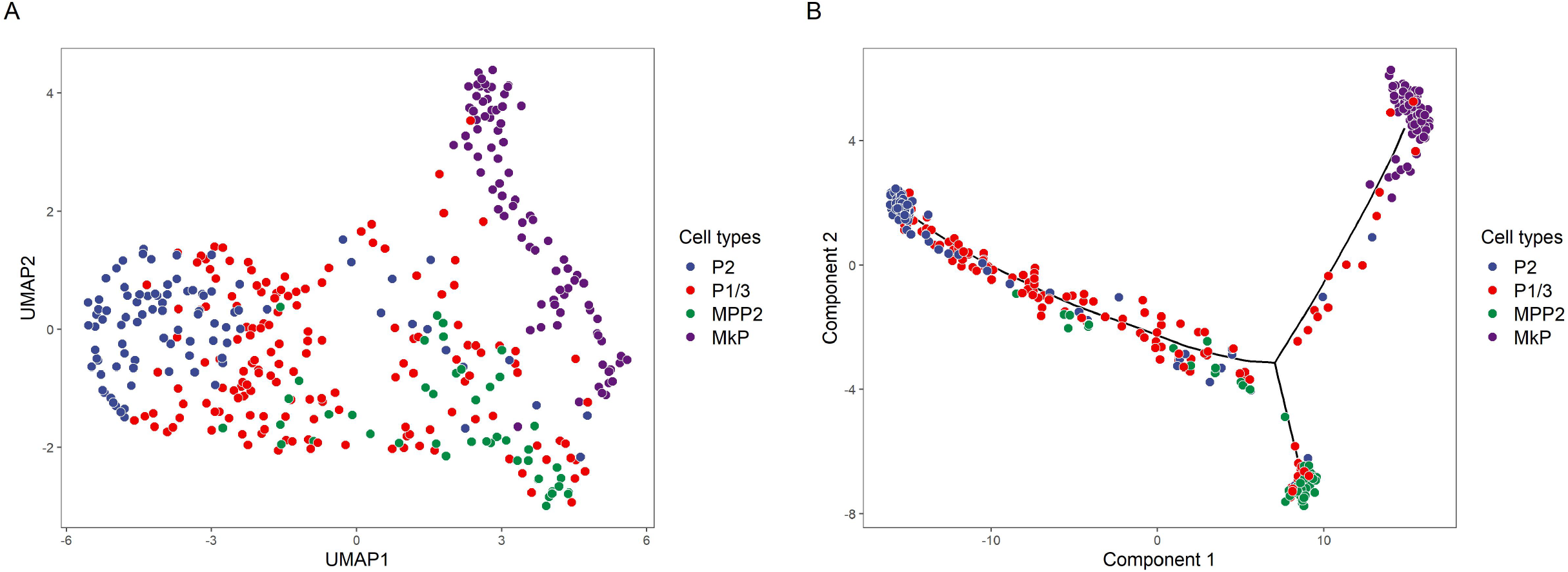
Cluster and trajectory analyses of single-cell RNA-Seq data. (A) UMAP plots of clusters. (B) Monocle trajectory. HSC1/HPC1-P2 cells are shown as P2. HSC1/HPC1-P1/3 cells are shown as P1/3. MPP2 and MkP cells were also analyzed. Data were obtained from analysis of 96 P2 cells, 180 from P1/3, 48 MPP2 cells, and 85 MkP cells.

Cluster analysis showed that HSC1/HPC1-P2 and MkP cells, respectively, form distinct clusters. In contrast, HSC1/HPC1-P1/3 and MPP2 cells were widely distributed between HSC1/HPC1-P2 and MPP2 clusters (Fig. 5A). Trajectory analysis suggested that HSC1/HPC1-P1/3 cells continuously go to the direction of MkP. Interestingly, MPP2 cells shared the common pathway with HSC1/HPC1-P1/3 cells until a certain point and then changed the direction (Fig. 5B). These data suggested that HSC1/HPC1-P1/3 cells lie between HSC1/HPC1-P2 and MkP cells along the differentiation, and MPP2 cells partially share the differentiation pathway with HSC1/HPC1-P1/3 cells.

We also analyzed differentially expressed genes in the HSC1/HPC1-P1/2/3 and MkP populations. The representative data are shown in Supplemental Fig. 4. The expression of *Vwf, Pl4, Cavin4*, and *Self* in the MkP population was significantly greater than that in the HSC1/HPC1-P1/2/3 populations, suggesting that HSC1/HPC1-P1/2/3 cells differ from MkP cells. The expression of *Proc, Mecom*, and *Sh2b3* in the HSC1/HPC1-P2 populations was significantly greater than that in the HSC1/HPC1-P1/3 and MkP populations. The expression of these genes in the HSC1/HPC1-P1/3 population was also significantly greater than that in the MkP population. The expression of *Runx1* in the MkP population was significantly greater than that in the HSC1/HPC1-P1/2/3 populations. Its expression in the HSC1/HPC1-P1/3 populations was also greater than that in the HSC1/HPC1-P2 populations. Taken together, these data supported that HSC1/HPC1-P1/3 cells transcriptionally lie between HSC1/HPC1-P2 and MkP cells.

### Mk gene expression in single cells

In general, single-cell RT-PCR is more sensitive than single-cell RNA-seq. Focusing on Mk- and HSC-related genes, the expression of 48 genes was selectively examined with 24 single cells each from HSC1/HSC2/HPC1-P1/2/3. Data are shown as a heatmap format in Supplemental Fig. 5.

*c-Kit* expression was detected in most cells from all populations (Supplemental Fig. 6). *Mpl* expression was also detected in most cells, except in HSC2-P3 cells. We detected Mk lineage markers (*CD41, CD61, Vwf*, and *Pf4*) as graphically shown in Fig. 6. The expression of *CD41* was detected in all HSC1/HSC2/HPC1-P1/2/3 populations although the frequencies of positive cells differed. *CD41*^+^*CD61*^+^ cells were detected in the HPC1-P1/2/3 populations (19 / 72 cells) slightly more than in the HSC1-P1/2/3 populations (10 / 72 cells). Very few *CD41*^+^*CD61*^+^ cells were detected in the HSC2-P1/2/3 populations. *Vwf* ^+^ cells were detected in the HSC1-P2 and HPC1-P1/2 populations whereas no *Vwf* ^+^ cells were detected in the HSC2-P1/2/3 and HSC1-P3 populations. A few *Vwf* ^+^ cells were detected in the HSC1-P1 and HPC1-P3 populations. The HPC1-P1/2 populations were significantly enriched in *CD41*^+^*Vwf* ^+^ cells, suggesting that the coexpression of *CD41/Vwf* may mark a certain stage of megakaryopoiesis. The HSC1-P1/3 and HPC1-P1/3 populations were significantly enriched in *CD41*^+^/*Pf4*^+^ cells, suggesting that the coexpression of *CD41/Pf4* is to some extent related to the Mk progenitor activity.

**Figure 6.**
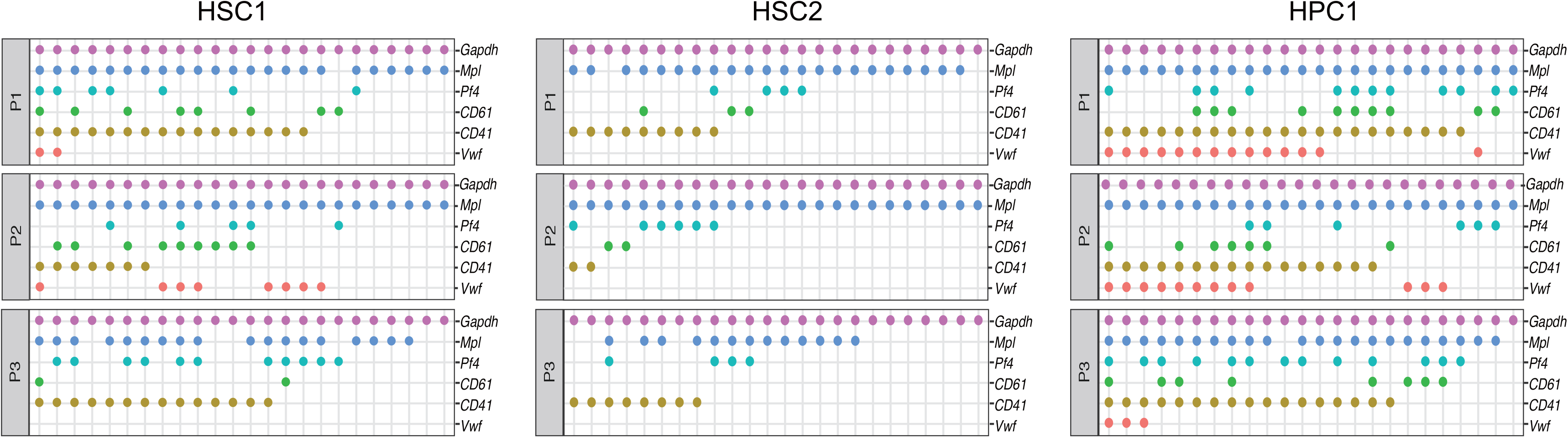
scRT-PCR of *Vwf, CD41, CD61, Pf4 and Mpl* genes. Each horizontal row represents 24 single cells. Positive cells defined as cells with a threshold cycle value (Ct) < 27.65 are shown as dots. *CD41*^+^*Vwf* ^+^ cells in HPC1-P1/2 (22 / 48 cells) were significantly more than those in the other populations (6 / 168) (p < 0.0001). *CD41*^+^/*Pf4*^+^ cells in HSC1-P1/3 and HPC1-P1/3 (33 / 96 cells) were significantly more than those in the other populations (7 / 120) (p < 0.0001) (Fisher’s exact test).

We also examined the regulators of Mks (Supplemental Fig. 6). *Nfe2*^+^, *Mef2a*^+^, and *Mef2c*^+^ cells were similarly detected in HSC1/HSC2/HPC1-P1/2/3. More *Gfi-1b*^+^ cells were detected in HSC1/HPC1-P1/2/3 and HSC2-P1 than HSC2-P2/3. HSC1-P3 and HPC1-P1/3 were relatively enriched in *Gata-1*^+^ cells. The relative expression of *GATA-1* was highest in HSC1-P3. *Gata2* expression was detected in all populations, but the frequency of positive cells was relatively low in HSC2-P3. Unpredictably, *Gata3* expression was detected in all populations (Supplemental Fig. 6). Taken together, these data suggested that the HSC1/HPC1-P1/3 populations were heterogeneous in Mk gene expression.

## Discussion

The HSC1/HSC2/HPC1-P2 populations were enriched in repopulating cells, as well as in multilineage colony-forming cells (Figs. 2, 3, 4). These data suggested that repopulating cells form multilineage colonies *in vitro*. Although the HSC1-P1/3 and HPC1-P1/3 populations barely repopulated in vivo, they appeared to be enriched in Mk colony-forming cells (Fig. 4, Supplemental Fig. 3). We named these early Mk-lineage committed progenitors MgPs. MgPs do not phenotypically overlap MkPs as summarized in Supplemental Table 4. MgPs partially overlap pMKPs which have been recently reported to be expanded in the mouse essential thrombocythemia model ^33^. MgPs should be more immature than MkPs ^12,14^ because MgPs generated significantly more Mks than did MkPs in vitro (Fig. 4D). To our knowledge, MgPs are the earliest Mk progenitors in the bone marrow of adult mice.

MgPs formed Mk colonies containing 8 cells on average, suggesting that single cells divided on average 3 times before endomitosis. To illustrate the relationship between MgPs and HSCs, we performed single-cell colony assay on 2-8 cells generated from single cells in culture (Supplemental Table 3). The loss of EM differentiation potential likely took place as early as from the first division. It should require 4 or more divisions to generate MgPs. These data suggested that MgPs are not immediate progeny of HSCs.

Single-cell colony assay of MPP2 cells revealed their multilineage and Mk potentials (Supplemental Fig. 3A). Notably, nmEMk colonies generated by MPP2 cells were significantly smaller than those generated by HSC1/HSC2/HPC1-P2 cells (Fig. 4D). Mk colonies generated by MPP2 cells were significantly smaller than those generated by MgPs, and Mk colonies generated by MkPs were significantly smaller than those generated by MPP2 (Fig. 4E). These data suggested that MkPs in the MPP2 population are downstream of MgPs, and MkPs in the Lin^-^c-Kit^+^Sca-1^-^CD150^+^CD41^+^ population are downstream of MkPs in MPP2 (Fig. 4E, Supplemental Fig. 3A) ^12^.

The expression of *CD41* and *Vwf* was together detected in the HPC1-P1/2 population, suggesting their expression may be related to a subset of MgPs (Fig. 6). Bioinformatics and functional assays have suggested that cells express both CD41 and Vwf when cells commit to the Mk lineage ^8,16^. Our data partially supported their claim. Notably, the combination of *CD41* and *Pf4* expression may be more likely to specify MgPs (Fig. 6).

Plt-RCs were detected in the HSC1-P2, HSC2-P2/3, and HPC1-P2 populations at very low frequencies (Fig. 3). The repopulating activity of Plt-RCs seemed to be lower, and their lifespan seemed to be shorter than those of Plt-biased HSCs ^8,9^. *Vwf* expression was detected in the HSC1/HPC1-P2 but not in HSC2-P2/3 populations. The expression of *Vwf* seemed irrelevant to Plt-repopulating activity (Fig. 6). Since we used 5×10^5^ instead of 2×10^5^ bone marrow cells as competitive cells, our transplantation assay might not be sensitive enough to detect Plt-biased HSCs with robust and stable Plt-reconstitution potential. Alternatively, the frequencies of Plt-biased HSCs might be lower than reported ^8-10^. HSCs likely give rise to MgPs at their early differentiation stage. To study how HSCs commit to Mk lineage, MgPs should serve as a good model. The molecular basis of lineage-biased HSCs is not known. This study suggested that lineage-bias fundamentally differs from lineage commitment in HSCs because MgPs differ from Plt-biased HSCs.

Based on our transplantation data, colony assay data, and trajectory analysis (Figs. 2-5), we assumed that the HSC1/HPC1-P2 populations give rise to the MPP2 and HSC1/HPC1-P1/3 populations. These populations give rise to the MkP population (Fig. 7A). The corresponding model for a hierarchical organization of LT-HSCs, CMPs, MgK, and MkPs is shown in Fig. 7B. MgP appeared to be a relatively homogenous population in terms of Mk colony-forming potential (Fig. 4, Supplemental Fig. 3). However, MgP turned out to be an extremely heterogeneous population in terms of gene expression profiles (Fig. 5). Data of single-cell RT-PCR supported the heterogeneity of MgPs (Fig. 6). We were unable to explain a discrepancy between functional and transcriptional data. Recent human studies have suggested that lineage commitment is a continuous process ^34-36^. If this were the case, there might be multiple differentiation and maturation steps from HSCs to MkPs through MgPs. In our model, Meg lineage commitment occurs in a stochastic manner at any differentiation stage between HSCs and CMPs. Further work is required to identify intermediate cells between LT-HSCs and MgPs along myeloid differentiation.

**Figure 7.**
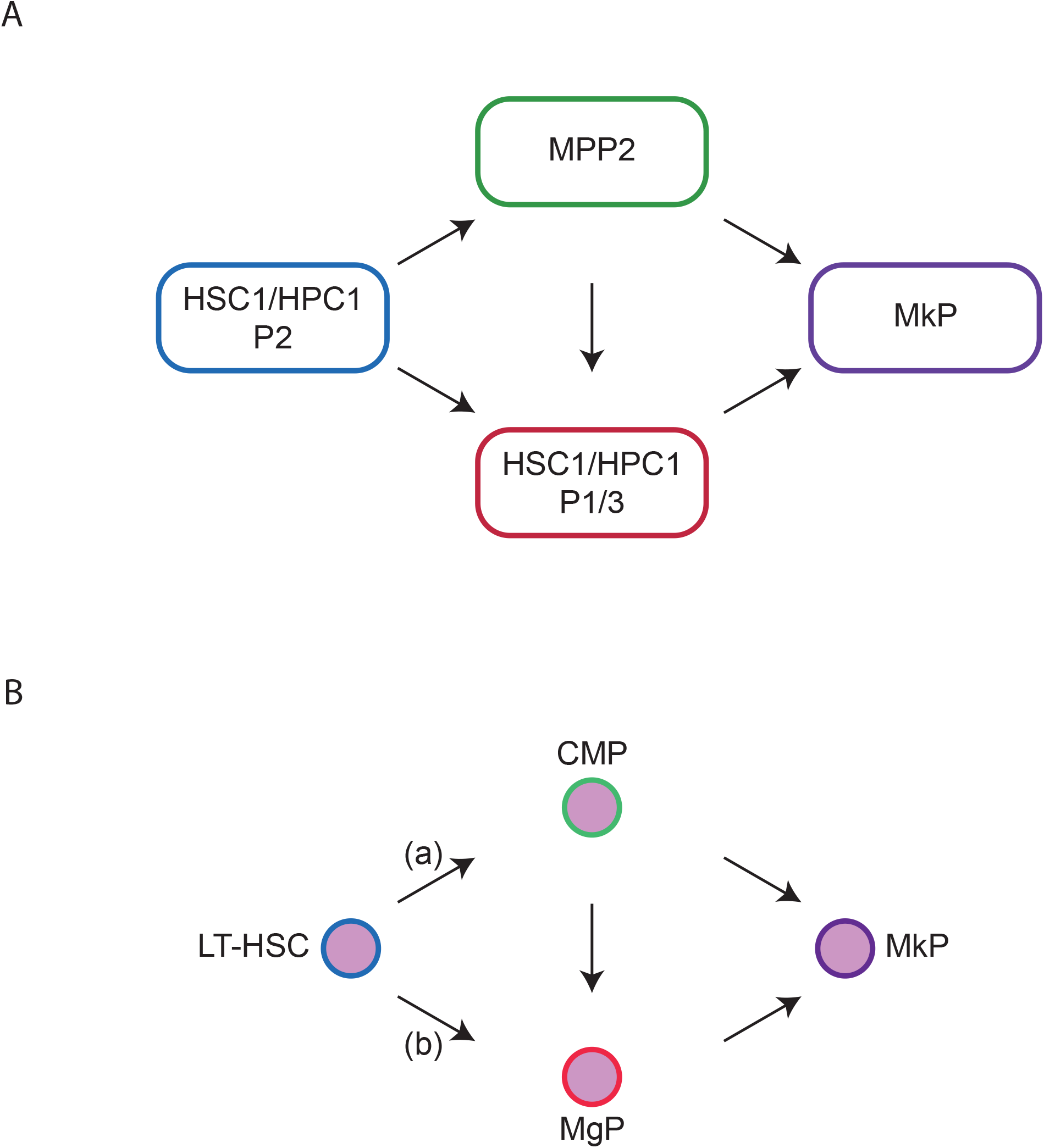
Hierarchical organization of HSCs and HPCs along Mk differentiation. (A) Population relationship. The HSC1/HPC1-P2 populations give rise to the MPP2 and HSC1/HPC1-P1/3 populations which then give rise to the MkP population. (B) Hierarchical relation among LT-HSCs, CMPs, MgPs, and MkPs. (a) LT-HSCs give rise to CMPs which then give rise to MgPs and MkPs. MgPs give rise to MkPs. (b) LT-HSCs give rise to MgPs as previously suggested ^37^.

### Database

scRNA-seq data were deposited at Gene Expression Omnibus with the accession number of GSE135013.

## Acknowledgements

This work was supported by grants from the National Key Research and Development Program of China Stem Cell and Translational Research (2017YFA0104900, 2019YFA0110203), CAMS Initiative for Innovative Medicine (CAMS-I2M) (2017-I2M-1-015), CAMS Fundamental Research Funds for Central Research Institutes (2019PT320017), and the National Natural Science Foundation of China (81670105, 81970119, 82070112).

## Author contribution

M.H., H.E. supervised the study; Z.L., J.W., M.X., P.W., Y.M., S.Z., X.W., F.D., H.C. performed experiments; Z.L., P. Z., M. H., H.E analyzed and interpreted data; Z.L., H.E. wrote the manuscript.

## Figure legends

**Supplemental Table 1.**
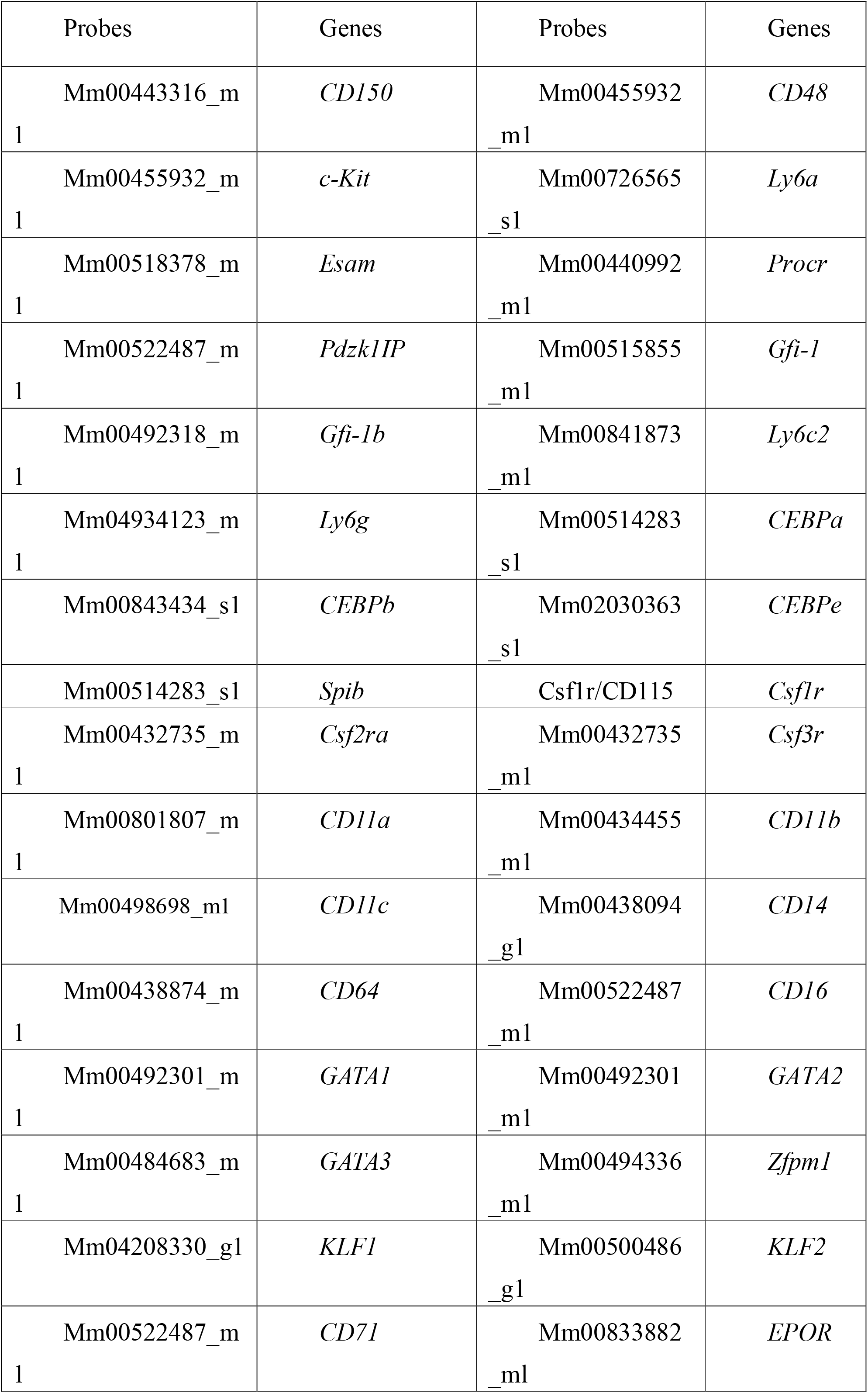
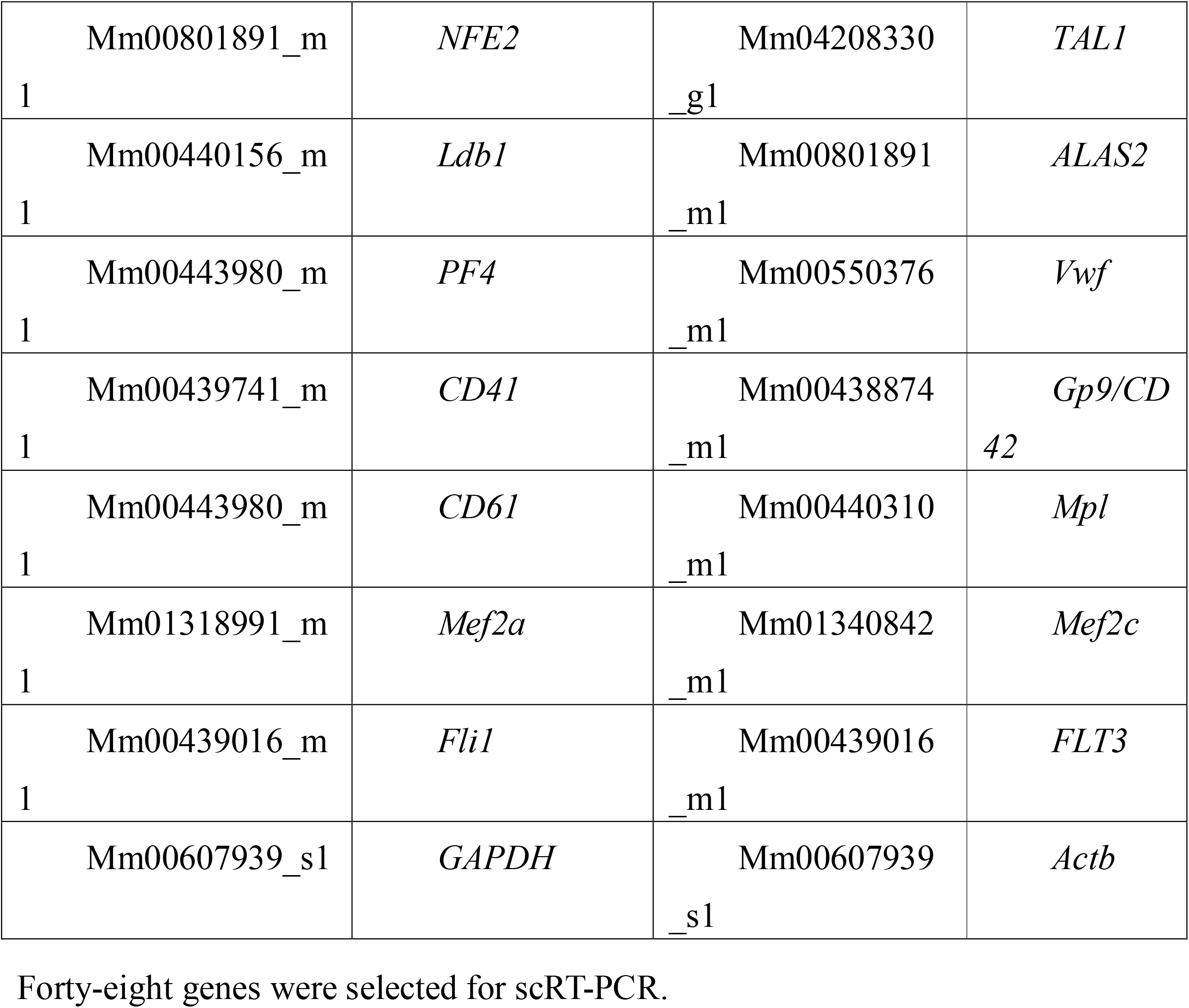
Gene set for scRT-PCR.

**Supplemental Table 2.** Paired daughter cells assay of HSC1/HPC1-P2 cells.

| Cells | One | The other | Pair No. |
| --- | --- | --- | --- |
| HSC1 | nmEMk | nmEMk | 7 |
|  | nmEMk | nmMk | 3 |
|  | nmEMk | nmE | 1 |
|  | nmEMk | nm | 4 |
|  | nmEMk | m | 3 |
|  | nmEMk | n | 1 |
|  | nmEMk | Mk | 1 |
|  | nmMk | nmE | 2 |
| HPC1 | nmEMk | nmEMk | 13 |
|  | nmEMk | nmMk | 5 |
|  | nmEMk | nm | 8 |
|  | nmEMk | n | 1 |
|  | nmEMk | m | 1 |

Single HSC1-P2 and HPC1-P2 cells were sorting and cultured in the medium of single-cell colony assay for 2 days. After single cells underwent the first division, paired daughter cells were individually transferred into new wells by micromanipulation and further incubated for 14 days. Colony cells were morphologically identified as neutrophils (n), macrophages (m), erythroblasts (E), and megakaryocytes (Mk). Paired daughter cells of which parental cells were assumed to have had the nmEMk differentiation potential are listed in the table.

**Supplemental Table 3.**
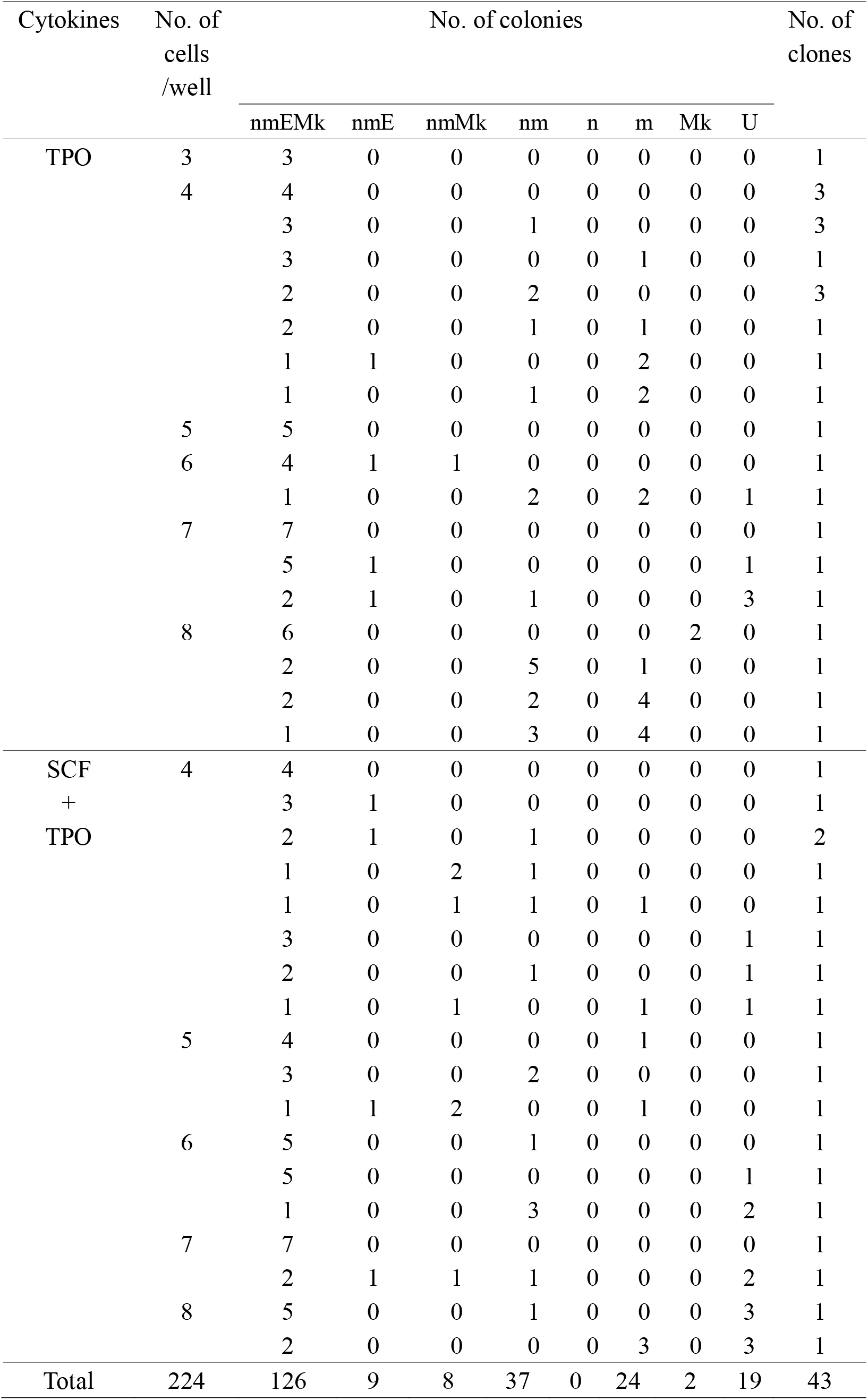
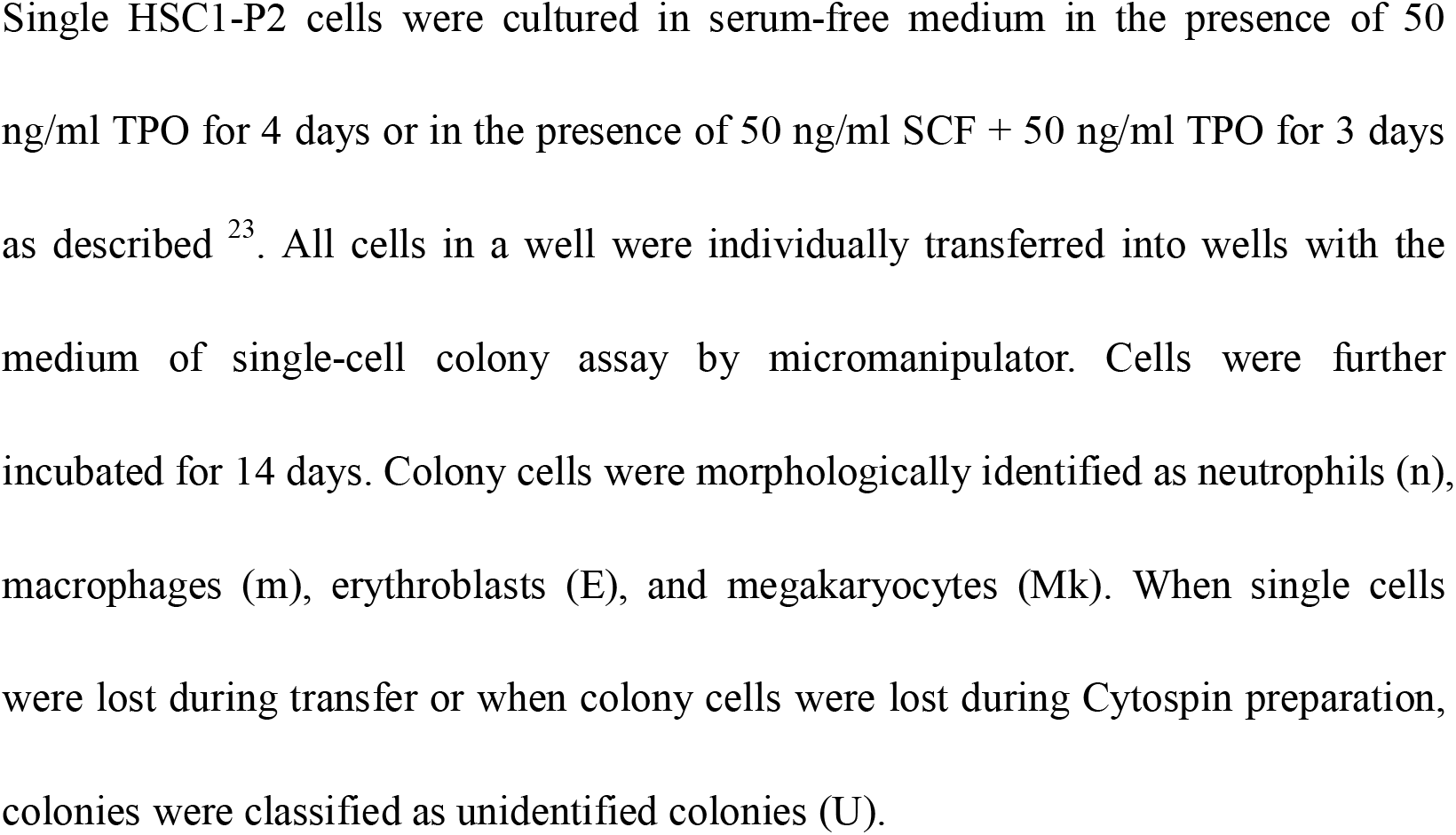
Myeloid lineage commitment in early divisions of HSCs.

**Supplemental Table 4.** Comparison of cell surface makers among HSC, MgP, and MkP populations.

| Cells<br>Populations | HSCs/CMPs |  | MgPs |  | MkPs |  |
| --- | --- | --- | --- | --- | --- | --- |
|  | HSC1 | HPC1 | HSC1/HPC1 | HSC1/HPC1 | MPP2 | MkP |
|  | P2 | P2 | P1 | P3 | (Passegue) | (Bryder) |
| CD201 | + | + | - | -/+ | -/+ | - |
| CD150 | + | + | + | + | + | + |
| CD48 | - | - | - | + | + | + |
| CD41 | - | + | -/+ | -/+ | -/+ | + |
| CD34 | - | - | - | - | + | + |
| c-Kit | + | + | + | + | + | + |
| Sca-1 | + | + | + | + | + | - |
| Lineage | - | - | - | - | - | - |
| CD135 | - | - | - | - | - | - |

“MkPs or MkP cells” were used as the name of functional megakaryocyte progenitors whereas “MkP” was used as the name of megakaryocyte progenitor population in this study. \*, MPP2 has been originally defined as a CD34-positive population. But, the CD34 marker was not used in Passegue’s study. -/+, both negative and positive cells are detectable.

## Supplemental figure legends

**Supplemental Figure 1. MPP2 and MkP gating** (A) The gating strategy of MPP2 cell is shown. The expression of CD34, CD41, and CD201 in MPP2 cells is additionally demonstrated. (B) The gating strategy of MkP cells is shown.

**Supplemental Figure 2. Morphological analysis of colony cells.** (A) Morphology of colony cells. Mk, megakaryocytes; n, neutrophils; m, macrophages; E, erythroblasts. (B) Different types of colonies: nmEMk, nmMk, nm, and m colonies are shown. Cells were collected from a well of 96-plates and spin down onto glass slides, and subjected to Wright Giemsa staining.

**Supplemental Figure 3. Additional colony assays.** (A) Single-cell colony assay for MPP2 and MkP. Sixty single cells were sorted and cultured for 14 days (n=3). The colony-forming efficiency of CD150^+^CD48^+^Flk2^-^c-Kit^+^Sca-1^+^Lin^-^ MPP2 cells was 52.2 ± 0.3 % (left panel). Most CD150^+^CD41^+^c-Kit^+^Sca-1^-^Lin^-^ MkP cells (98.9%) only gave rise to Mks (right panel). (B) Thirty single cells from the HSC1/HSC2/HPC1-P1/2/3 populations were cultured for 21 days in two independent experiments. On day 21, colonies were scored, and cell components were identified. The frequency of nmEMk colony-forming cells in HSC1-P2 cells was significantly greater than those in HSC1-P1 cells (71.1% versus 29.6%, p < 0.001) and HSC1-P3 cells (71.1% versus 30.6%, p < 0.001). The frequency of nmEMk colony-forming cells in HSC2-P2 cells was significantly greater than that in HSC2-P3 cells (39.1% versus 14.3%, p < 0.001). The frequency of nmEMk colony-forming cells in HPC1-P2 cells was significantly greater than those in HPC1-P1 cells (58.6% versus 13.8%, p < 0.001) and HPC1-P3 cells (58.6% versus 7.1%, P < 0.001) by chi-square test. The frequency of Mk colony-forming cells in HSC1-P2 cells was significantly lower than those in HSC1-P1 cells (4.4% versus 40.7%, p < 0.001) and HSC1-P3 cells (4.4% versus 50.0%, p < 0.001). The frequency of Mk colony-forming cells in HPC1-P2 cells was significantly lower than those in HPC1-P1 cells (24.1% versus 65.5%, p < 0.001) and HPC1-P3 cells (24.1% versus 78.6%, p < 0.001) by chi-square test.

**Supplemental Figure 4. Gene expression analysis of scRNA-seq data. (A-D)** The expression of Mk-related genes (*Vwf, Pl4, Cavin2*, and *Selp*) and **(E-H)** the expression of HSC-related genes (*Procr, Runx1, Mecom*, and *Sh2b3*) are selectively shown after a standard log normalization step. *, p<0.05; **, p<0.01; ***, p<0.001; ****, p<0.0001 (Wilcoxon test).

**Supplemental Figure 5. Single-cell RT-PCR analysis of HSC1/HSC2/HPC1-P1/2/3.** The heatmaps with 48 genes in the horizontal rows and 24 single cells in the vertical columns. Positive cells are defined as the threshold cycle values [Ct] < 27.65.

**Supplemental Figure 6. Single-cell RT-PCR analysis of Mk-related gene.** Twenty-four single cells were sorted from the HSC1/HSC2/HPC1-P1/2/3 populations by flow cytometry. RT-PCR was performed on these single cells to compare the expression of the 48 genes. The positive cell ratios of selected genes are shown in the histograms (left panels). Positive cells were defined as cells with a threshold cycle value (Ct) < 27.65 are. The relative expression levels for the positive cells are shown in the violin plots (right panels). The relative expression level was defined as (28-Ct) values.

