## Supplemental Figs. 1-6 for "Early megakaryocyte lineage-committed progenitors in adult mouse bone marrow"

A

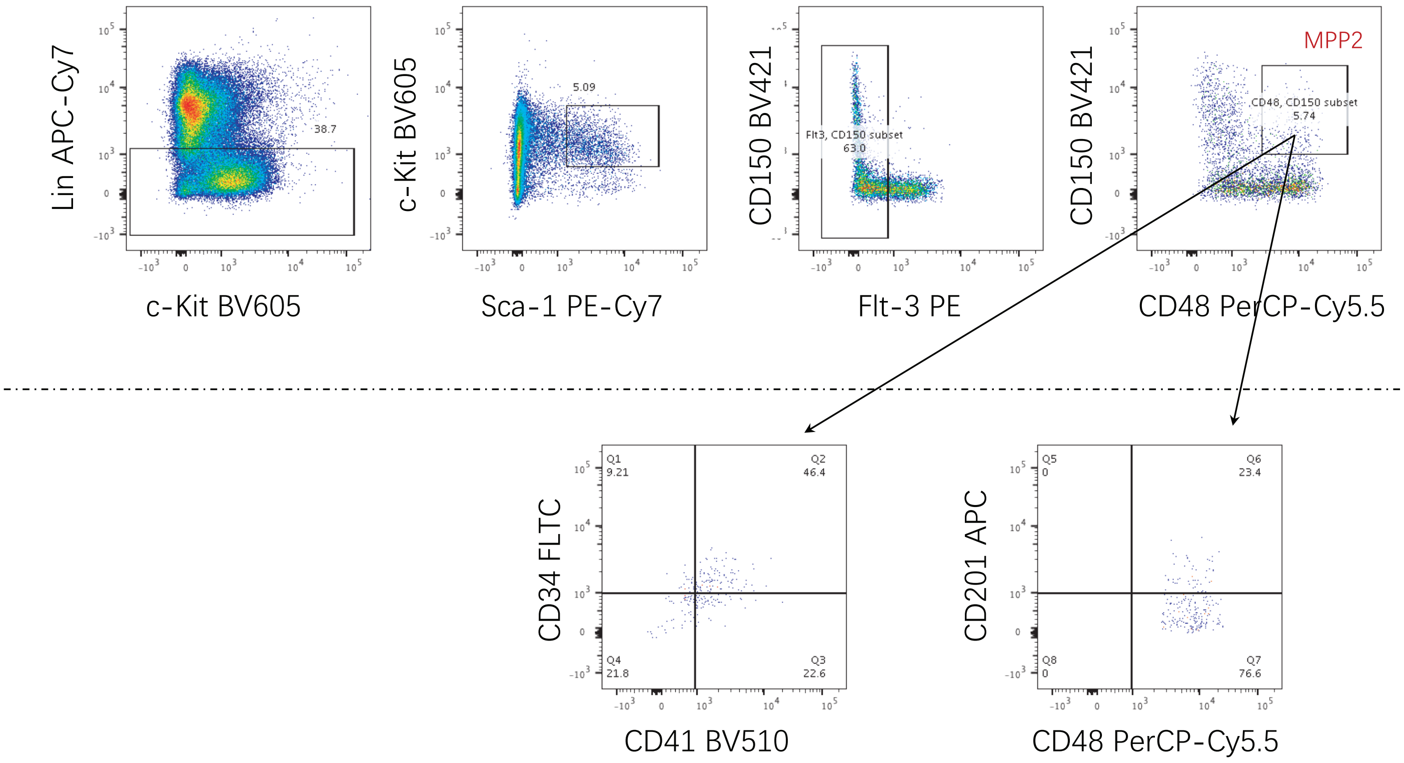


B

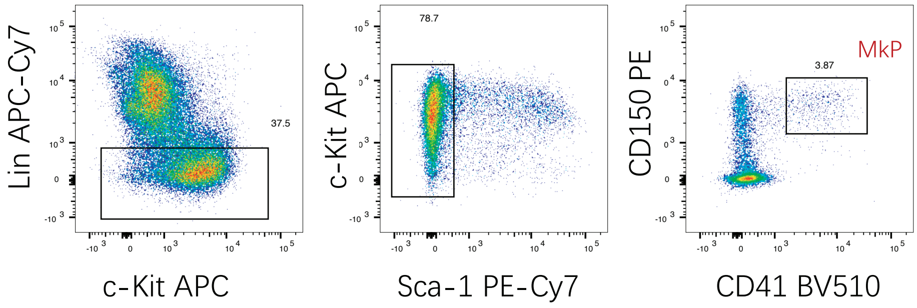


ZL_Supplemental Figure 1

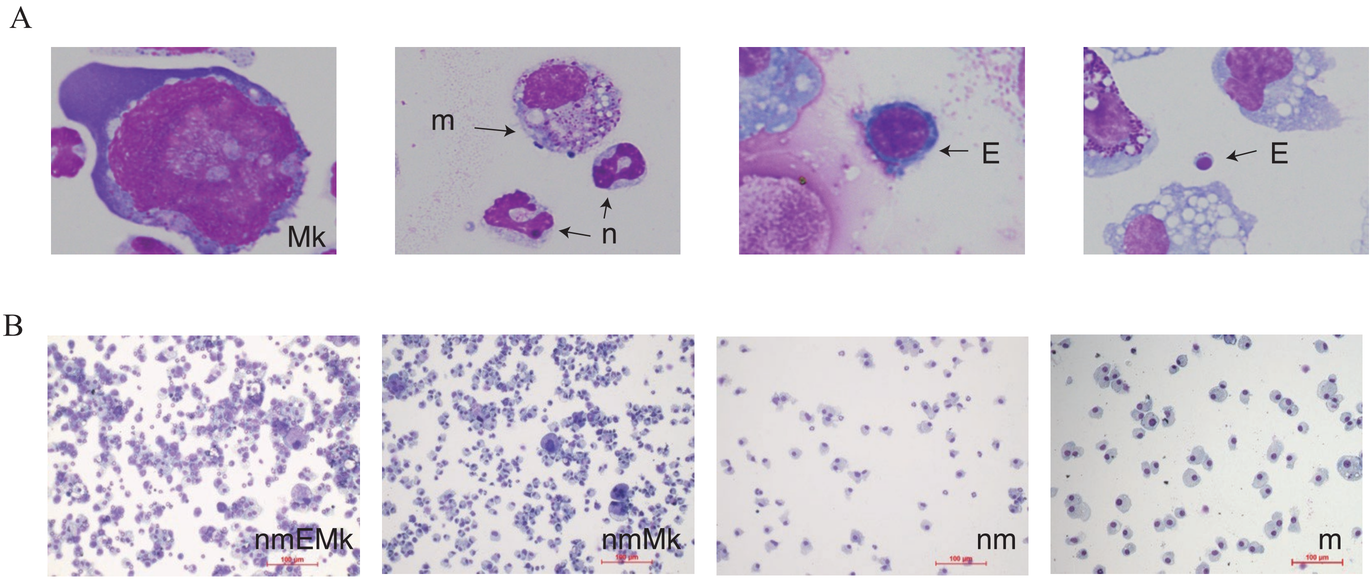


ZL_Supplemental Figure 2

A

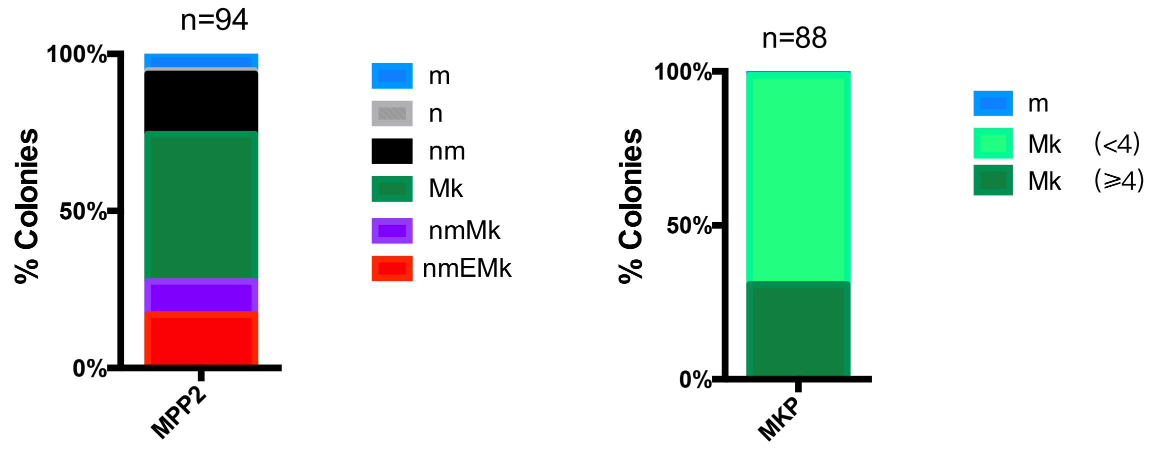


B

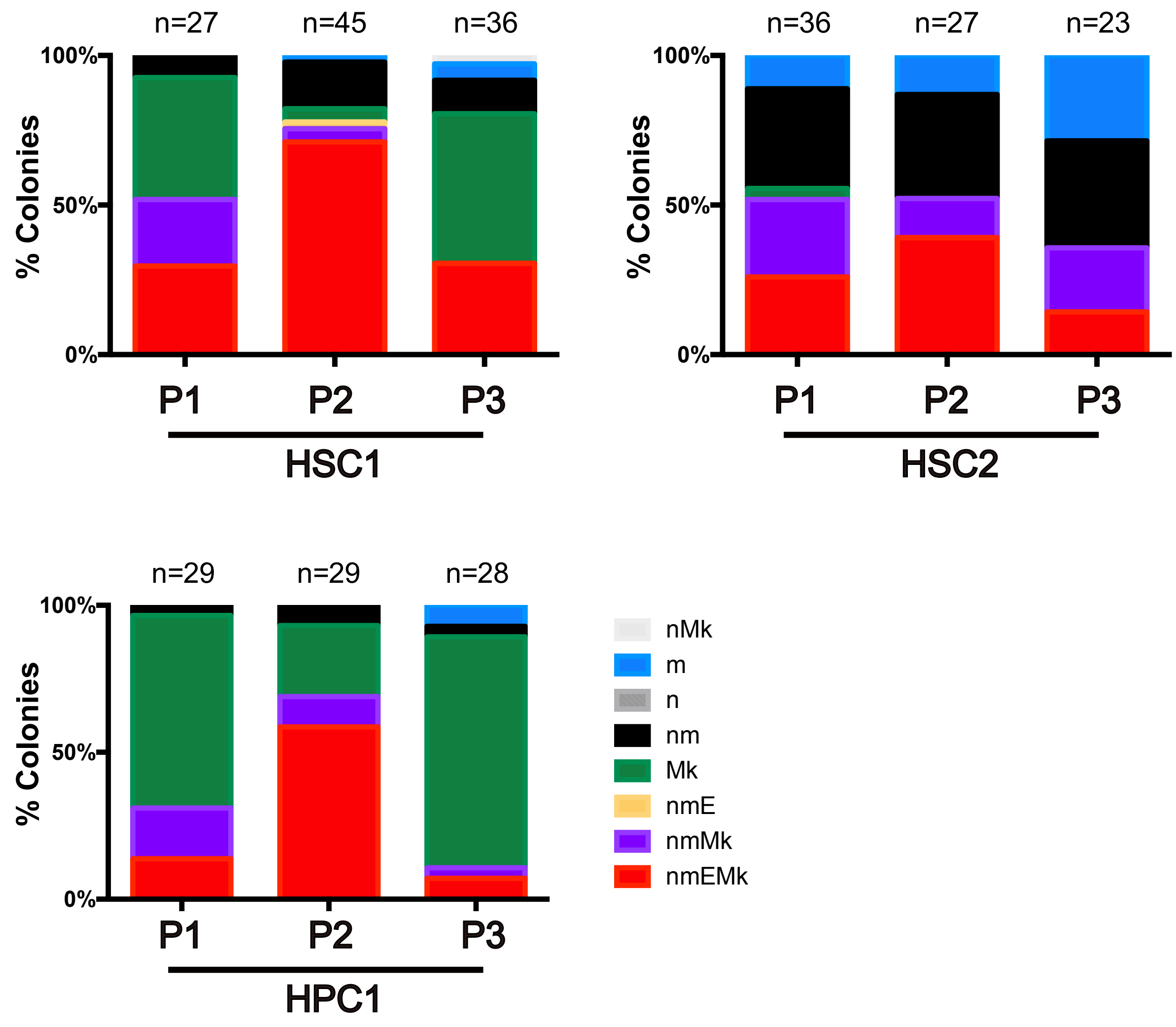


ZL_Supplemental Figure 3

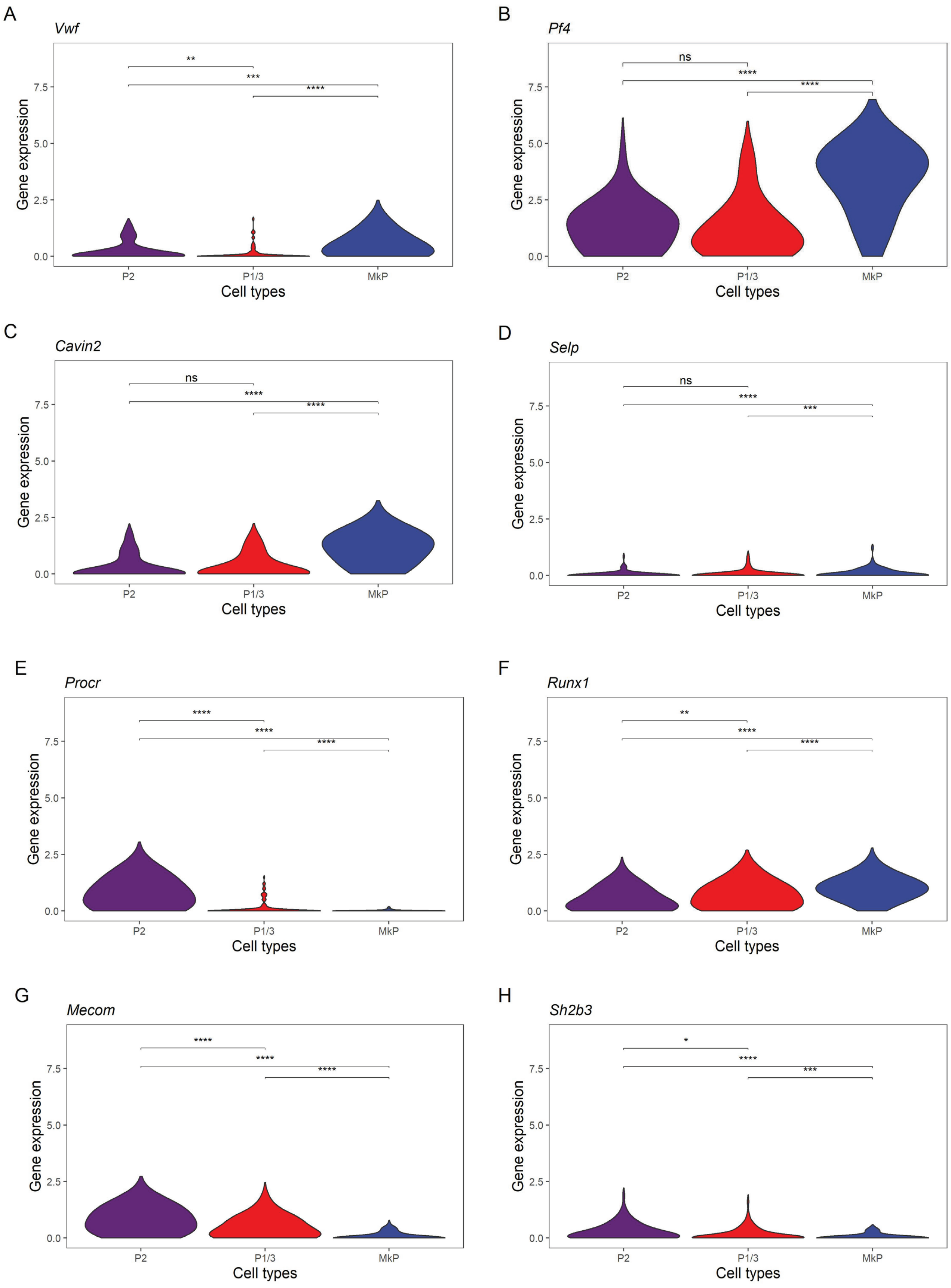


ZL_Supplemental Figure 4

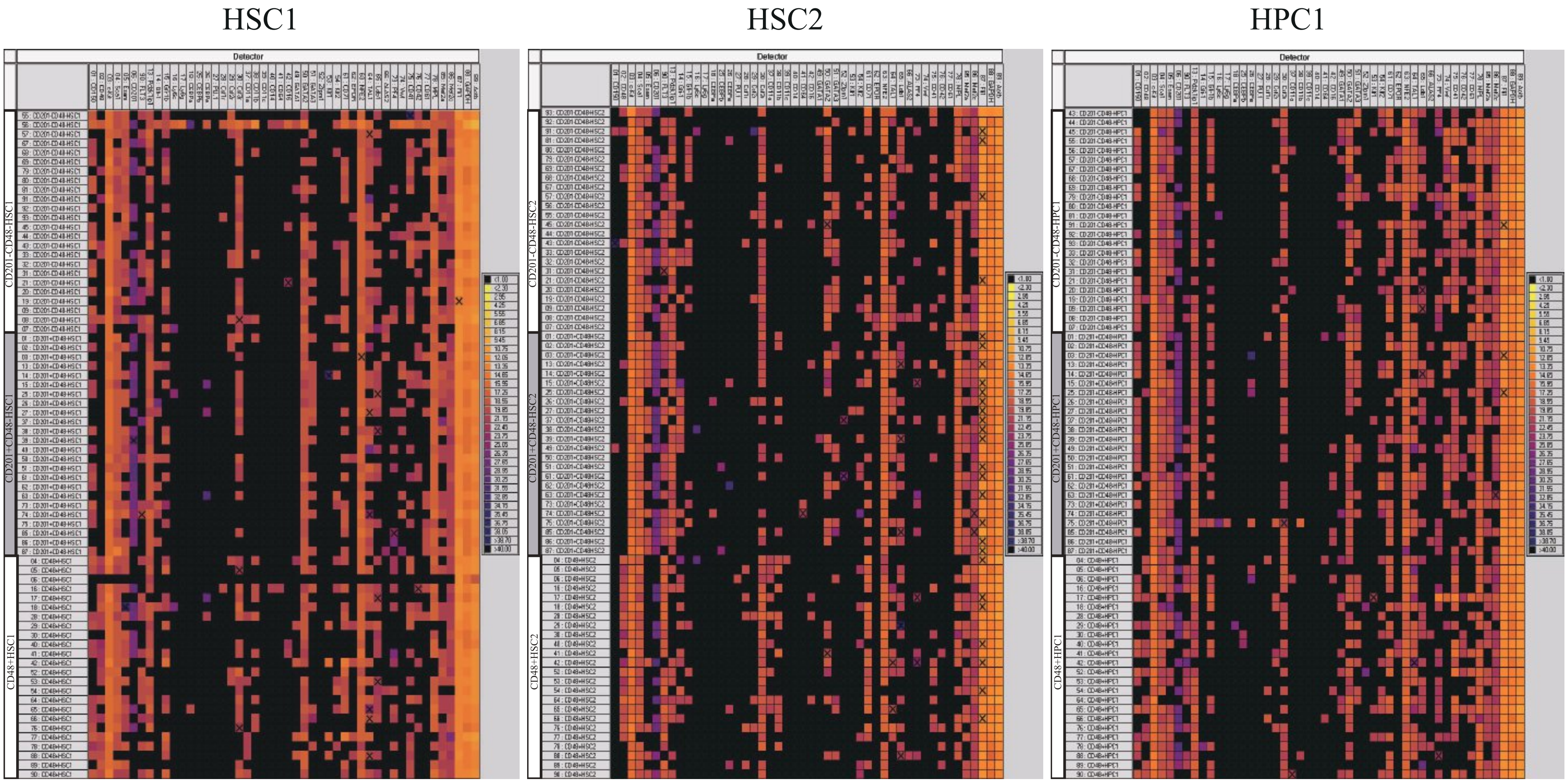


ZL_Supplemental Figure 5

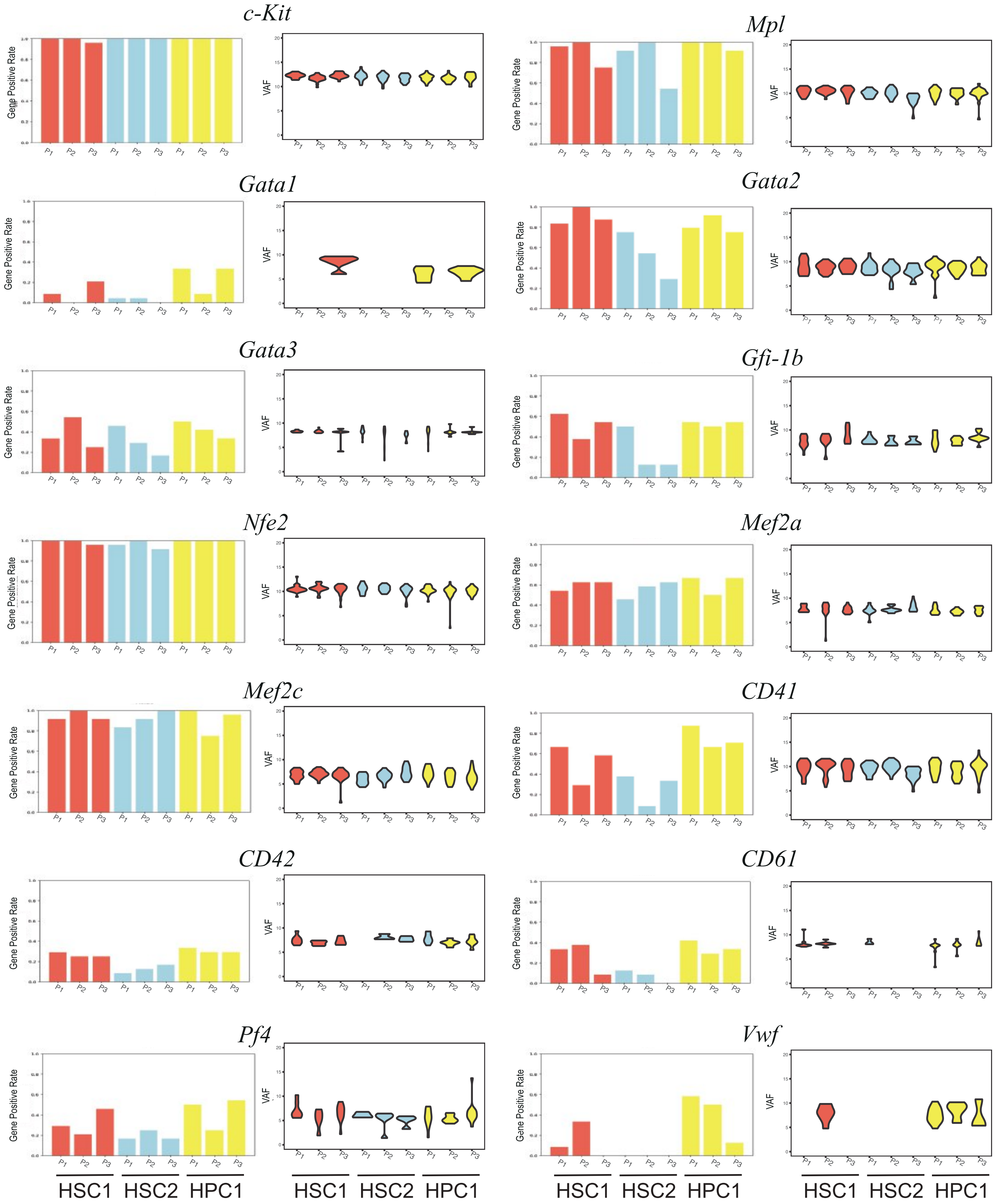


ZL_Supplemental Figure 6
